# Conditional Generative Modeling for De Novo Protein Design with Hierarchical Functions

**DOI:** 10.1101/2021.11.10.467885

**Authors:** Tim Kucera, Matteo Togninalli, Laetitia Meng-Papaxanthos

**Author notes:** These authors jointly supervised the work (Matteo Togninalli, Laetitia Meng-Papaxanthos).

## Abstract

**Motivation:** Protein design has become increasingly important for medical and biotechnological applications. Because of the complex mechanisms underlying protein formation, the creation of a novel protein requires tedious and time-consuming computational or experimental protocols. At the same time, machine learning has enabled the solving of complex problems by leveraging large amounts of available data, more recently with great improvements on the domain of generative modeling. Yet, generative models have mainly been applied to specific sub-problems of protein design.

**Results:** Here we approach the problem of general purpose protein design conditioned on functional labels of the hierarchical Gene Ontology. Since a canonical way to evaluate generative models in this domain is missing, we devise an evaluation scheme of several biologically and statistically inspired metrics. We then develop the conditional generative adversarial network ProteoGAN and show that it outperforms several classic and more recent deep learning baselines for protein sequence generation. We further give insights into the model by analysing hyperparameters and ablation baselines. Lastly, we hypothesize that a functionally conditional model could generate proteins with novel functions by combining labels and provide first steps into this direction of research.

**Availability:** Code and data is available at https://github.com/timkucera/proteogan

## 1 Introduction

Designing new proteins with a target biological function is a frequent task in biotechnology and has broad applications in synthetic biology and pharmaceutical research, for example in drug discovery (Huang *et al*., 2016). The task is challenging because the sequence-structure-function relationship of proteins is extremely complex and not yet fully understood (Dill and MacCallum, 2012). Protein design is therefore mostly done by trial-and-error methods, such as directed evolution (Arnold, 1998), which rely on a few random mutations of known proteins and selective pressure to explore a space of related proteins. This process can be time-consuming and cost-intensive, and most often only explores a small portion of the sequence space. At the same time, data characterizing proteins and their functions is readily available and constitutes a promising opportunity for machine learning applications in protein sequence design.

Multiple generative models have recently been proposed to design proteins for different tasks, such as developing new therapies (Mueller *et al*., 2018; Davidsen *et al*., 2019), enzymes (Repecka *et al*., 2021), nanobody sequences (Riesselman *et al*., 2019; Shin *et al*., 2021) or proteins that lead to antibiotic resistance (Chhibbar and Joshi, 2019). These models are typically focused on a sub-task of protein design and thus are limited to a given application, often even to a specific protein family. This requires retraining for a new task, which limits the diversity and number of sequences from which a model can learn. In other domains, such as the closely related natural language generation, one can observe a trend towards general-purpose models that are then used in various contexts (Brown *et al*., 2020). We posit that, also in protein design, a one-fits-all model may learn common underlying principles across different protein classes improving the quality of generated sequences. Going further, it may even be able to create not only novel sequences, but novel functions by combining different aspects of functionality it has learned in different protein families.

We therefore develop ProteoGAN, a general purpose generative model for conditional protein design based on the Molecular Function Gene Ontology (GO), a hierarchy of labels describing aspects of protein function. These functions vary from binding specific agents, to transporter or sensor activity, catalysis of biochemical reactions and many more. Here, additionally, the information encoded in the hierarchical organisation may help model performance. We base our model on the popular Generative Adversarial Network (GAN) framework because of their recent success on the generation of enzymes that are soluble and display catalytic activity when they are experimentally tested (Repecka *et al*., 2021). We extend the framework by proposing a conditional mechanism to incorporate the multilabel hierarchical information of protein function into the generation process.

However, developing such a generative model can be challenging, not least because problem-specific evaluations are lacking. An evaluation metric needs to assess whether a generated sample is valid (i.e. realistic and functional), a hard problem in itself, and further needs to be fast to compute on a large number of samples. The evaluation of generative models is still ongoing research, particularly in the domain of protein design (Papineni *et al*., 2002; Salimans *et al*., 2016; Heusel *et al*., 2017; Shmelkov *et al*., 2018; Kynkäänniemi *et al*., 2019; DeVries *et al*., 2019). While gold-standard validation of a generated sequence implies the synthesis of the proteins in the lab, the lack of *in-silico* assessments makes it difficult to efficiently compare methods for protein sequence design.

We therefore compose an array of evaluation metrics for generative protein design based on the Maximum Mean Discrepancy (MMD) statistic to measure distributional similarity and conditional consistency of generated sequences with real proteins. We further propose measures to account for sequence diversity.

### 1.1 Related generative models for protein design

#### 1.1.1 Guided and conditional protein generative models

Machine learning models and more recently deep generative models (Eddy, 2004; Goodfellow *et al*., 2014; Kingma and Welling, 2014; Rezende *et al*., 2014; Vaswani *et al*., 2017; Li *et al*., 2017) have been used to design *in silico* biological sequences, such as RNA, DNA or protein sequences (Durbin *et al*., 1998; Davidsen *et al*., 2019; Brookes *et al*., 2019), often with the aim to create sequences with desired properties. There are two main strategies to achieve this, one is guided and the other conditional. Guided approaches use a predictor (also called oracle) in order to guide the design towards target properties, through iterative training-generation-prediction steps (Brookes *et al*., 2019; Gane *et al*., 2019; Angermueller *et al*., 2019; Gupta and Zou, 2019; Killoran *et al*., 2017; Repecka *et al*., 2021; Gligorijevic *et al*., 2021). In a scenario with multiple functional labels however, the lack of highly accurate and fast multilabel predictors for protein function impairs guided-generation techniques in functional protein generation (Zhou *et al*., 2019). Conditional approaches integrate the functional information in the generation mechanism itself, eliminating the need for a predictor. For example, Madani *et al*. (2020) developed ProGen, a conditional transformer that enables a controlled generation of a large range of functional proteins, but the need for a sequence context can be experimentally constraining and is not compatible with *de novo* design. Ingraham *et al*. (2019) present a graph-based conditional generative model that relies on structural information, which is only sparsely available. Das *et al*. (2018) and Greener *et al*. (2018) train conditional VAEs in order to generate specific proteins, such as metalloproteins. Karimi *et al*. (2019) used a guided conditional Wasserstein-GAN to generate proteins with novel folds. All these models either focus on a sub-task of protein design only, or rely on context information such as 3D structure or template sequence fragments. Here we propose a general purpose model for protein design that only requires specifying the desired functional properties for generation.

#### 1.1.2 Evaluation of generative models

To this date, there is no definitive consensus on the best evaluation measures for the evaluation of *quality*, *diversity* and *conditional consistency* of the output of a (conditional) generative model (Papineni *et al*., 2002; Salimans *et al*., 2016; Heusel *et al*., 2017; Shmelkov *et al*., 2018; Kynkäänniemi *et al*., 2019; DeVries *et al*., 2019). Most measures that stand out in computer vision such as the Inception Score (IS) (Salimans *et al*., 2016), the Frechet Inception Distance (FID) (Heusel *et al*., 2017), or GAN-train and GAN-test (Shmelkov *et al*., 2018) depend on an external, domain-specific predictor. For functional protein design such predictors are neither good nor fast enough to entirely rely on their predictions when evaluating and training neural networks. The Critical Assessment for Functional Annotation (CAFA) challenge reports the currently best model (NetGO) with an F_max_ score of 0.63, which has a prediction speed of roughly 1000 sequence per hour (F_max_ is the maximal F1-score over confidence thresholds, see their paper or our supplemental section 1) (Radivojac *et al*., 2013; Zhou *et al*., 2019; You *et al*., 2019). On the contrary, the domain-agnostic duality gap can be computed during training and at test time, and has been shown to correlate well with FID (Grnarova *et al*., 2019).

In natural language modeling, perplexity is a common evaluation metric which relates to the probability of a test set under the model. This however requires access to a likelihood which is not available in some models, suchasGANs, and is not always a good indicator of sample quality (Theis *et al*., 2016). Another approach measures how many wild-type residues can be recovered from an incomplete sequence, which however goes against the idea of de novo protein design.

Despite the increasing interest of the research community for protein generation models, no clear metrics have emerged as reliable tools to compare them.

## 2 System and methods

### 2.1 Evaluation framework for conditional protein sequence design

Generative models are difficult to evaluate because there is no ground truth one could compare each generated sample with. Instead, the goal of generative modeling is to create data that is similar in its properties but not identical to some target data. Evaluation is further complicated when the data can not be straightforwardly validated, such as in protein design where a generated sample would need to be physically synthesized to prove its functionality. We therefore propose to assess the quality of a model by comparing its generated sequences to natural protein data, with principled statistical tests.

We compose an array of metrics to evaluate conditional generative models for protein design which are based on two-sample goodness-of-fit statistics that can be computed for structured data such as protein sequences and the resulting high-dimensional feature vectors. They have the advantage to be fast to compute and to be differentiable, and can therefore be used during training, for hyperparameter selection, early stopping or as a loss. We corroborate these metrics by comparing the statistics computed with biologically relevant embeddings.

The following sections detail specific aspects of the evaluation and the respective metric we devised.

#### 2.1.1 Evaluating *distribution similarity* with MMD

As generative models aim to model the distribution of target data, it is a natural choice to evaluate them with a statistical two-sample test that compares generated and training data distributions. This approach is difficult to apply to protein sequence data directly but can be applied to extracted feature vectors. We propose to use Maximum Mean Discrepancy (MMD) (Gretton *et al*., 2012), a test statistic that compares mean embeddings in a Reproducing Kernel Hilbert Space (RKHS). In the past, MMD has been used to infer biological pathways or sequence homology from biological sequences or for distinguishing sets of structured biological sequences (Leslie *et al*., 2001; Vegas *et al*.,2016; Borgwardt *et al*., 2006).

Let 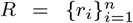 and 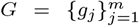 be samples from the distributions of real and generated proteins sequences, respectively *P_r_* and *P_g_*. Then:

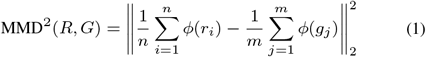

We decided to use the (normalized) Spectrum kernel (Leslie *et al*., 2001) with its feature map *ϕ* since it is fast to compute and sufficiently complex to distinguish protein properties of interest, which we validate in meta evaluations of the metrics in the Discussion section. The feature mapping counts the occurrences of kmers in a sequence (*k* = 3, resulting in 8000 features). We verify that our measure is robust to the choice of kernel by using an alternative Gaussian kernel (supplemental Section 11).

To confirm our evaluations with the Spectrum kernel feature map we further compute MMD using the biological embeddings ProFET (Ofer and Linial, 2015), a collection of handcrafted features, UniRep (Alley *et al*., 2019), a learned protein embedding based on an LSTM recurrent network and ESM-1b, a learned protein embedding based on a transformer language model (Rives *et al*., 2021). We remove kmer related features in ProFET to avoid confounding with our Spectrum kernel based metrics, resulting in ca. 500 features which were then scaled to the same range. UniRep has a dimensionality of 1900, and we use the mean hidden state over the sequence as the representation. The ESM embedding has 1280 features, where we use the mean hidden representation of the 33rd layer.

We further confirm the results computed by the MMD statistic with a second statistic, the Kolmogorov-Smirnov (KS) test, which is more expensive to compute (supplemental Section 13).

#### 2.1.2 Evaluating *conditional consistency* with MRR

For conditional generation specifically, we need to assess the model’s capability to generate sequences consistent with some target labels. We extend the MMD metric by computing MMD between subsets of sequences for each label and ranking the RKHS distance between generated samples and their target label among distances to off-target labels. It measures how many sets of real sequences with off-target labels are closer in distribution to the generated sequences than real sequences with the target label.

Let *R* be a set of real sequences *R_i_* with annotated labels *i* ∈ {1, …, *d*}, where *d* is the total number of labels. Let 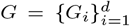 be an equally structured set of generated sequences. We want to maximise the following mean reciprocal rank (MRR):

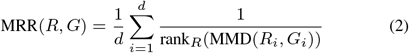

where rank_*R*_(MMD(*R_i_, G_i_*)) is the rank of MMD(*R_i_, G_i_*) among elements of the sorted list [MMD(*R*_1_, *G_i_*), MMD(*R*_2_, *G_i_*), …, MMD(*R_d_, G_i_*)]. MRR(*R, G*) is maximal and of value 1 when the generated distributions of proteins for a given label are the closest to the distribution of real proteins with the same label.

Since the GO protein function labels we are using are organized in a hierarchy we also include a variant of MRR that gives more insight on conditional performance for closely-related functions in the label hierarchy, by not penalizing ranking errors arising from parent and children labels (*MRR_B_*).

#### 2.1.3 Evaluating the *diversity* of generated sequences

A common failure mode of generative models and specifically in GANs is mode collapse, where a model produces a single mode of the data (Salimans *et al*., 2016). In practice, we would like to ensure diversity of generated samples in order to represent a significant part of the space of possible sequences while ensuring that the sequences remain realistic. In order to capture this trade-off we consider three measures. First, we monitor the duality gap (Grnarova *et al*., 2019) of our GAN model (Figure S7). A small duality gap indicates good convergence and common failure modes, such as mode collapse, can be detected. Second, we propose two heuristic *diversity estimates* of the distributions of generated and real sequences. These are the average entropy over feature dimensions (*n* = 1000 bins) as well as the average pairwise RKHS distance between sequences. Ideally, we would expect these two statistics in the generated data to exhibit small differences (noted ΔEntropy or ΔDist.) relative to the real distribution (i.e. as measured in the test set). Finally, to ensure that we are not overfitting to the training data, we also report nearest-neighbour squared euclidean distances over the spectrum kernel embeddings for our model between the feature maps of the generated sequences and training sequences, and control that they are not closer than to the testset (Figure S8).

#### 2.1.4 A note on out-of-distribution evaluation

A particularly interesting aspect of generative protein modeling is the creation of novel sequences. While this is already useful for in-distribution samples which expand the repertoire of existing proteins with new variants, an exciting outlook is the generation of out-of-distribution (OOD) data which would correspond to a novel kind of protein. The evaluation of OOD generation is however notoriously difficult (Ren *et al*., 2019; Nalisnick *et al*., 2019). We go first steps into this direction by holding out five manually selected label combinations from the training data and generating sequences conditioned on these label combinations after training. We then report Top-*X* accuracy (*X* ∈ 1, 10) where a generated sequence is counted accurate if a true sequence from the held out sample is among its *X* nearest neighbors in embedding space. The OOD sets contain approximately 1000 sequences each and the number of real sequences that are not in the OOD sets is a multiple of the number of sequences in the OOD set, with a multiplication factor varying from 2 to 30. The held out label names and GO identifiers can be found in supplemental Table S2.

While this metric should give a sense of OOD generation capability in a comparison of different models, we note that biological plausibility of such truly novel OOD samples remains to be shown.

### 2.2 A conditional generative model for hierarchical multi-label protein design

After having set the framework to evaluate and compare models for generative protein design, we now develop a conditional generative model to generate proteins with desired functions. While most existing *de novo* protein sequence generation models focus on a specific function, we here aim at modeling different biological functions at the same time. Hence, we introduce ProteoGAN, a conditional GAN (cGAN) able to generate protein sequences given a large set of functions in the Gene Ontology (GO). The GO is a curated set of labels describing protein function and is organized in a directed acyclic graph. We are therefore dealing with a hierarchical multilabel problem.

We explore several conditioning mechanisms, label embeddings and model architectures to find well-suited configurations specifically for hierarchical multilabel protein design. The final model is found by an extensive hyperparameter search guided by our metrics MMD and MRR. We then analyse the results of the optimization by functional Analysis of Variance (fANOVA (Hutter *et al*., 2014)) to give insights about model parameters. The following sections detail conditioning mechanisms and variants thereof which we propose, the general model architecture and the hyperparameter optimization scheme.

#### 2.2.1 Model architecture

We focus on GANs due to their promising results on protein sequence design tasks (Repecka *et al*., 2021). Our base model is a Wasserstein-GAN with Gradient Penalty (Arjovsky *et al*., 2017; Gulrajani *et al*., 2017). It contains convolutional layers and skip connections in both the generator and the discriminator, its funnel-like structure is similar to DCGAN (Radford *et al*., 2015). In the generator, the label is concatenated to the latent noise vector input of the network. In the discriminator we explore various mechanisms for conditioning. Exact model configurations are determined through a hyperparameter search detailed in section 6 (Figure 1).

**Fig. 1:**
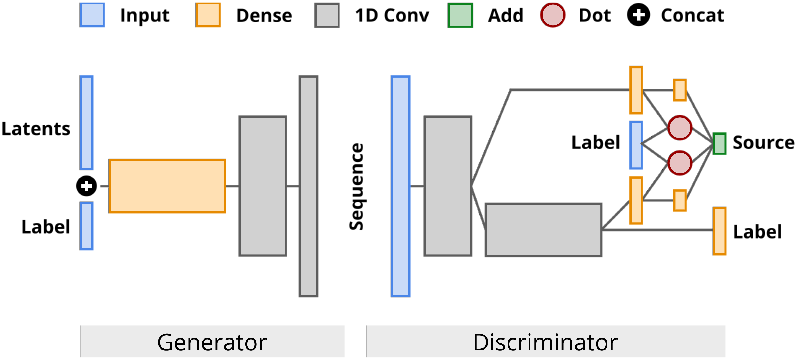
Architecture of ProteoGAN after extensive hyperparameter optimization. We varied the number of layers, conditioning mechanism(s), number of projections, convolutional filters, and label embeddings, among others.

#### 2.2.2 Conditioning

We assess the performance of three types of conditioning mechanisms during our hyperparameter search: projection(s), auxiliary classifiers, or a combination of both. To the extent of our knowledge, there is no generative model that uses either multiple projections or a combination of projections and auxiliary classifiers in the literature.

In the conditional GAN with *projection* discriminator, as introduced in Miyato and Koyama (2018), the discriminator *D* is decomposed into a sum of two terms, one being the inner product between a label embedding and an intermediate transformation of the sequence input, and the second term being solely dependent on the sequence input. Let (***x***, ***y***) ~ *p* be a sample from the dataset, where ***x*** is a one-hot encoded sequence, and ***y*** an encoding of choice of the categorical label. The projection discriminator can be written as 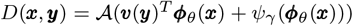, with ***y*** → ***ν***(***y***) a linear projection of the label encoding, ***φ**_θ_* an embedding function applied to the sequence ***x***, *Ψ_γ_* a scalar function applied to the embedding function ***φ**_θ_* and 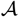 an activation function of choice.

We propose an extension to the projection mechanism, namely use *multiple projections* in the discriminator. We hypothesize that this could help utilizing protein sequence information at the different abstraction levels of the convolutional layers. In addition to the previous notations introduced in this section, let us assume that we have now *k* projections. Let 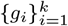 be *k* neural networks, which can be decomposed in *n_i_* layers 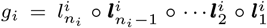. Let 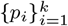 be the layer number at which the inner product with the output of the linear projection 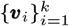 occurs in each neural network. The projections obey a tree-like branching structure, where all layers *p* ≤ *p_i_* of the neural network *i* are shared with the neural networks *j* for which *p_i_* < *p_j_*, and the branching of a different projection is always done at a different layer number. The discriminator with multiple projections can then be written as 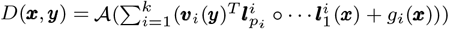.

We further propose to include an *auxiliary classifier* (AC) (Odena *et al*., 2017) *C_D_* to the objective function in addition to the projections, combining two previously independently used conditioning mechanisms. The AC shares parameters with the discriminator except a label classification output layer and adds a classification loss term to both the generator and discriminator: 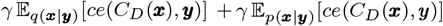, where ce denotes the cross-entropy function, ***x*** ~ *q*(***x**|**y***) the learned conditional distribution and **x** ~ p(**x**|**y**) the conditional distribution of the data. *C_D_* is trained together with the discriminator loss and predicts the labels of the real or generated sequences.

The conditioning mechanisms are further explained in the supplementary Section 2.

#### 2.2.3 Hierarchical label encoding and physicochemical amino acid properties

Given the hierarchical structure of the functional labels we allow for three types of label encodings ***y***: (i) one-hot encoding, as a common encoding for labels, (ii) Poincaré encoding (O’neill, 2014), which embeds labels in a hyperbolic space that is well-suited for hierarchical data and (iii) node2vec encoding (Grover and Leskovec, 2016), which preserves neighbourhood relationships by encoding the nodes of the GO directed acyclic graph (DAG) based on random walks. All of these encodings capture the relations between labels in the GO DAG and that way incorporate the information into the GAN. If a protein has multiple GO labels, the label encodings are summed to represent the set of assigned GO labels. We further allow to concatenate physicochemical properties of the respective amino acids to the encoding of the sequences. These are obtained from the AAIndex (Kawashima *et al*., 2007) (https://www.genome.jp/aaindex), a list with accession numbers can be found in Table S1.

## 3 Implementation

### 3.1 Data

Sequence data and GO labels were obtained from the UniProt Knowledgebase (UniProtKB, Consortium (2019)) and filtered for experimental evidence, at least one existing GO annotation, standard amino acids and a maximum length of 2048. It resulted in 157891 sequences total. We restricted functional labels to a total number of 50, imposing a minimum threshold of approximately 5000 sequences per label. In the GO DAG, sequences automatically inherit the labels of their parents, therefore such missing labels were imputed to complete the hierarchical information. We note that sequences exhibiting one of five manually selected label combinations (named A-E) were held out from the training data, to test the OOD performance of our model (see Section S2 for further details about the label combinations).

We randomly split the dataset in training, validation and test sets keeping ca. 15000 (10%) sequences in both the validation and test sets. During hyperparameter optimization, we use smaller splits with ca. 3000 sequences each. We ensure that all labels have a minimum amount of samples in the test and validation sets, and use the same number of sequences per class for the calculation of our MRR measure (1300 and 300 sequences, respectively). Further details about the dataset and splits are available in the supplementary Section 3 and Figure S1.

Since we do two-sample tests we do not remove homologous sequences from the test set, but for completeness we report our results with homology control up to 50% in supplemental section 15 (compare also the approach of Bileschi *et al*. (2019)).

### 3.2 Baseline comparisons

We compare our model ProteoGAN with classic probabilistic language models and more recent deep-learning models for protein sequence generation based on the Variational Autoencoder (VAE) and transformer framework:

- *HMM*: A profile HMM (Eddy, 2004) for each individual label (marked OpL for ‘one-per-label’) and for each label combination (marked OpC for ‘one-per-combination’). The former discards multilabel information and totals 50 models, the latter accounts for 1828 models, one model for each label combination present in the training set.
- *n-gram*: An n-gram model (*n* = 3) for each individual label (marked OpL, discards multi-label information, total of 50 models) and for each label combination (marked OpC, total of 1828 models).
- *CVAE*: A conditional Variational Autoencoder for proteins from Greener *et al*. (2018). We adjusted the model to incorporate the 50 labels of our problem setting and performed a Bayesian optimization hyperparameter search.
- *ProGen*: A language model from Madani *et al*. (2020) based on a state-of-the-art transformer architecture. Conditional information is included by prepending label tokens to the sequence. We reduced model size and retrained on our dataset.

We also perform ablation studies on ProteoGAN to understand the influence of several aspects of the model:

- *one-per-label GAN (OpL-GAN)*: One instance of ProteoGAN for every label, with the conditioning mechanism removed (total of 50 models). Sequences for a target label are generated by sampling them from the GAN trained on the sequences annotated with the same label. With this model, we assess whether training 50 models can be replaced by a conditioning mechanism.
- *Predictor-Guided*: ProteoGAN without conditioning mechanism, which results in a single GAN trained on the full data. The generated sequences are then annotated with the labels predicted by a state-of-the-art predictor NetGO (You *et al*., 2019). Comparing to this model allows us to investigate how a predictor model guiding the GAN compares to a conditional GAN.
- *Non-Hierarchical*: Same as ProteoGAN, but trained without the hierarchical multi-label information. Each sequence is included multiple times with each of its original labels separately. For fairness, we keep the number of gradient updates the same as for the other models. With this model we explore the usefulness of accounting for the GO hierarchy.

We refer the reader to the respective papers and to the supplementary Section 4 for further information on the baseline models.

### 3.3 Hyperparameter optimization

We conducted two Bayesian Optimization and HyperBand (BOHB) searches (Falkner *et al*., 2018) on six Nvidia GeForce GTX 1080, first a broad search among 23 hyperparameters (1000 models) and a second, smaller and more selective, among 9 selected hyperparameters (1000 models) on a maximum of 27 epochs. The optimization objective was set to maximize the ratio of our evaluation measures MRR/MMD to balance between distribution similarity and conditional consistency of the generated sequences. Both searches were complemented by a functional analysis of variance (fANOVA) (Hutter *et al*., 2014). The 27 best selected models of the second hyperparameter search were then trained for a prolonged duration of 100 epochs, the best performing model of these (i.e. ProteoGAN) then for 300 epochs. Further details about hyperparameter optimization are available in the supplementary Section 5.

## 4 Discussion

### 4.1 Meta-evaluation of metrics: Spectrum MMD is an efficient metric for protein design

Different embeddings capture different aspects of the original data. We were interested whether the relatively simple Spectrum kernel embedding would be sufficient to assess distribution similarity and conditional consistency, and hence compared it to three biologically founded embeddings: ProFET (Ofer and Linial, 2015), a handcrafted selection of sequence features relating mostly to biophysical properties of single amino acids or sequence motifs, UniRep (Alley *et al*., 2019), an LSTM-based learned embedding, and ESM (Rives *et al*., 2021), a transformer-based learned embedding. The latter two were shown to recover various aspects of proteins, including structural and functional properties as well as evolutionary context.

We compared these embeddings by scoring their ability to classify protein structure and function, for which we trained Support Vector Machines (SVM) trained on each of the embeddings. We classify domains of the CATH protein structure classification database (10000 sampled out of 500000, 10 repetitions, 80 – 20 train-test split), where we report balanced accuracy (Table 1), and we classify the 50 GO functional terms of our dataset (10000 sampled out of 150000, 10 repetitions, 80 – 20 traintest split), where we report the Fmax score used in the CAFA challenge, compare Zhou *et al*. (2019) and supplemental Section 1) (Table 1).

**Table 1.** Classification results of SVMs trained on different embeddings, for the structural classes of the CATH database (balanced accuracy), and for 50 functional classes of the GO (*F*_max_ score). All values in percent.

| Embedding | C | A | T | H | GO |
| --- | --- | --- | --- | --- | --- |
| Spectrum | 65 ± 3 | 54 ± 4 | 57 ± 2 | 47 ± 3 | 58 ± 1 |
| ProFET | 53 ± 2 | 38 ± 4 | 48 ± 2 | 28 ± 3 | 52 ± 1 |
| UniRep | 72 ± 4 | 73 ± 6 | 68 ± 2 | 58 ± 3 | 71 ± 1 |
| ESM | 77 ± 3 | 91 ± 1 | 86 ± 1 | 79 ± 2 | 80 ± 1 |

The ESM embedding is arguably the most powerful in this comparison and expectedly achieved the best scores. Notably though, the Spectrum kernel embedding is also considerably well suited to assess aspects of proteins on the structure and function, while being orders of magnitudes faster to compute and requiring less compute resources. This makes it more fit for the requirements on performance during evaluation or hyperparameter optimization of neural networks and other models. Another reason to choose the Spectrum kernel embedding is its simplicity, as it makes no assumption on the data distribution: The learned embeddings UniRep and ESM are complex non-linear mappings trained on large amounts of natural sequences, and while they perform great on natural in-distribution data, their behavior on generated sequences remains unpredictable. Moreover, when evaluating artificial sequences, embeddings are generally affected by the choice of parameters in the kernel as we show in the Supplemental Section 11. The Spectrum embedding has proven to be the most robust in this regard. We therefore propose it as the primary feature map in our evaluation metrics, and confirm it with evaluations based on the other embeddings (Tables S6,S7,S8). To validate MMD itself as a well-suited test statistic for protein design we confirm it with feature-wise KS statistics (Figures S13,S14,S15,S16).

### 4.2 Hyperparameter analysis: The conditional discriminator of ProteoGAN is most critical to its performance

We tested a wide range of hyperparameters and architectural choices for cGANs and analysed them in a fANOVA framework with respect to the protein design performance metrics MMD and MRR. To inform subsequent work on these models, we could empirically derive several design principles for GANs specifically for protein design (please refer to supplemental Section 6 for the raw marginal predictions of all parameters from which we deduce the following statements).

To begin with, smaller architectures performed much better than networks with more than four hidden layers. This size seems to be sufficient to model proteins, although of course the optimization places a selective pressure towards fast converging (small) models. The generator was less sensitive to its learning rate, while the discriminator showed strong preference towards learning rates below 1e-3. This may arise from the increased burden on the discriminator from the secondary training objective for classifying labels. It follows that it is more important that the discriminator arrives at an optimal solution rather than at local optima often found by larger learning rates.

We observed a trade-off between distribution similarity and conditional consistency. This manifested in increasing MRR and decreasing MMD performance when weighing stronger the training loss term of the auxiliary classifier, and also when switching between the different conditioning mechanisms. While we could not show significant impact of our proposed multiple projections, the combination of both conditioning mechanisms showed clear improvements in conditional performance.

We surprisingly found that a simple one-hot encoding of the labels showed the best results when comparing different label embeddings capturing the hierarchical relationships between labels. We also observe that only using the sequence as input, as opposed to appending biophysical feature vectors to the sequence embedding, led to the best performance. The amino acid identity, rather than its properties, appears to be more critical to sequence modeling.

Moreover, the discrete one-hot label embedding seems to be easier to interpret for the model than the continuous node2vec embeddings or the hyperbolic Poincaré embeddings. While these embeddings contain more information, the one-hot encoding presents them in a more accessible form. Also, hyperbolic spaces require special operators for many basic concepts that a neural network would need to learn first (Ganea *et al*., 2018).

Other popular extensions to the GAN framework such as input noise, label smoothing or training ratios did not significantly affect model performance in our context (compare Figures S5, S6). Summarizing, a small model with both conditioning mechanisms and no further sequence or label augmentation worked best. Further improvements to the architecture should focus on improving the discriminator, as hyperparameters affecting it showed the most impact (Figure S5). Our final model ProteoGAN is the best performing model of the optimization and has multiple projections, an auxiliary classifier, no biophysical features and one-hot encoding of label information.

### 4.3 Baseline comparisons: ProteoGAN outperforms other methods

Based on the proposed metrics for distribution similarity, conditional consistency and diversity we assess the performance of ProteoGAN and compare it to several baselines. The results are consolidated by an evaluation with the biological embeddings ProFET, UniRep and ESM, as well as feature-wise KS-statistics of the embeddings (Tables 2,S6,S7, Figures S13,S14,S15).

**Table 2.**
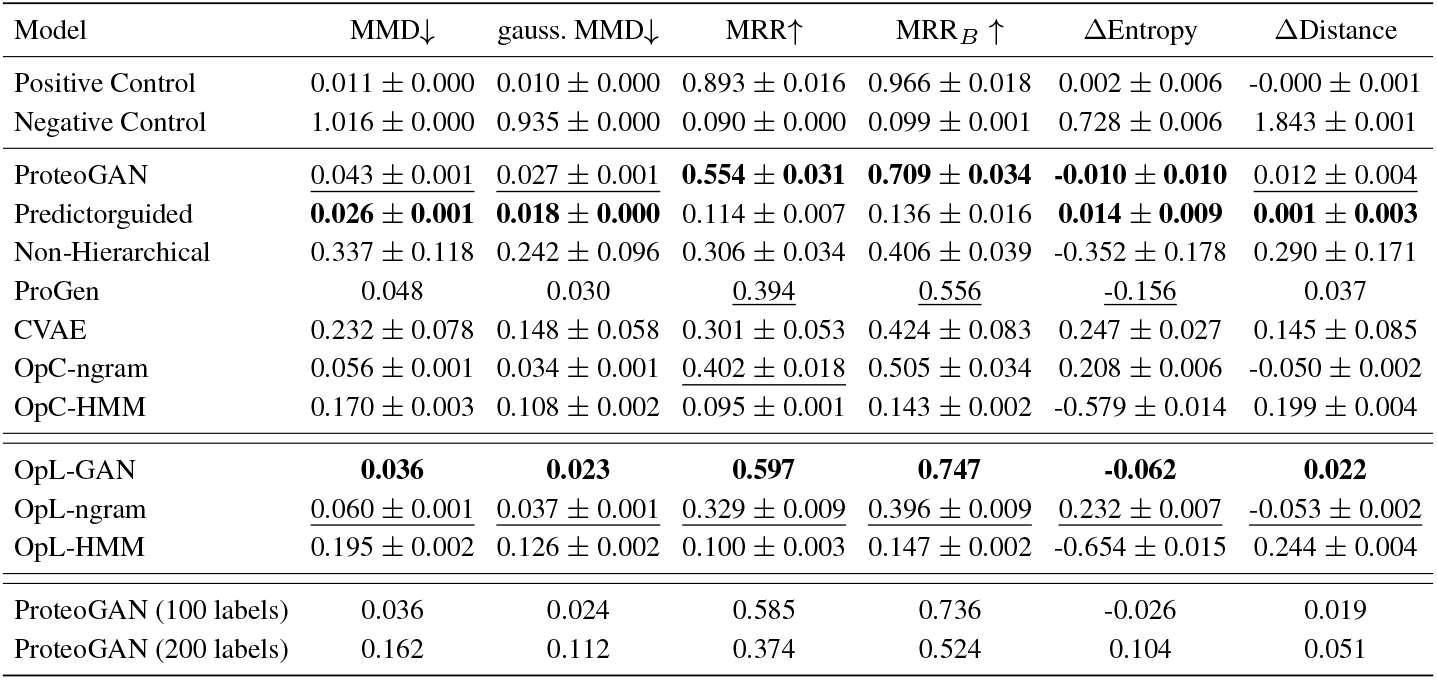
Evaluation of ProteoGAN and various baselines with MMD, MRR and diversity metrics based on the Spectrum kernel embedding (the results of other embeddings can be found in Table S6 and S7). An arrow indicates that lower (↓) or higher (↑) is better. The positive control is a sample of real sequences and simulates a perfect model, the negative control is a sample that simulates the worst possible model for each metric (constant sequence for MMD, randomized labels for MRR, repeated sequences for diversity measures). Best results in bold, second best underlined. Given are mean values and standard deviation over five data splits. Due to the computational effort, the OpL-GAN and ProGen were only trained on one split. The second and third panel show models presented in Section 3.2. The fourth panel shows variants of ProteoGAN that were trained on different datasets with more labels (100 or 200).

| Model | MMD↓ | gauss. MMD↓ | MRR↑ | MRR$_B$ ↑ | ΔEntropy | ΔDistance |
| --- | --- | --- | --- | --- | --- | --- |
| Positive Control | 0.011 ± 0.000 | 0.010 ± 0.000 | 0.893 ± 0.016 | 0.966 ± 0.018 | 0.002 ± 0.006 | -0.000 ± 0.001 |
| Negative Control | 1.016 ± 0.000 | 0.935 ± 0.000 | 0.090 ± 0.000 | 0.099 ± 0.001 | 0.728 ± 0.006 | 1.843 ± 0.001 |
| ProteoGAN | <u>0.043 ± 0.001</u> | <u>0.027 ± 0.001</u> | **0.554 ± 0.031** | **0.709 ± 0.034** | **-0.010 ± 0.010** | <u>0.012 ± 0.004</u> |
| Predictorguided | **0.026 ± 0.001** | **0.018 ± 0.000** | 0.114 ± 0.007 | 0.136 ± 0.016 | **0.014 ± 0.009** | **0.001 ± 0.003** |
| Non-Hierarchical | 0.337 ± 0.118 | 0.242 ± 0.096 | 0.306 ± 0.034 | 0.406 ± 0.039 | -0.352 ± 0.178 | 0.290 ± 0.171 |
| ProGen | 0.048 | 0.030 | <u>0.394</u> | <u>0.556</u> | <u>-0.156</u> | 0.037 |
| CVAE | 0.232 ± 0.078 | 0.148 ± 0.058 | 0.301 ± 0.053 | 0.424 ± 0.083 | 0.247 ± 0.027 | 0.145 ± 0.085 |
| OpC-ngram | 0.056 ± 0.001 | 0.034 ± 0.001 | <u>0.402 ± 0.018</u> | 0.505 ± 0.034 | 0.208 ± 0.006 | -0.050 ± 0.002 |
| OpC-HMM | 0.170 ± 0.003 | 0.108 ± 0.002 | 0.095 ± 0.001 | 0.143 ± 0.002 | -0.579 ± 0.014 | 0.199 ± 0.004 |
| OpL-GAN | **0.036** | **0.023** | **0.597** | **0.747** | **-0.062** | **0.022** |
| OpL-ngram | <u>0.060 ± 0.001</u> | <u>0.037 ± 0.001</u> | <u>0.329 ± 0.009</u> | <u>0.396 ± 0.009</u> | <u>0.232 ± 0.007</u> | <u>-0.053 ± 0.002</u> |
| OpL-HMM | 0.195 ± 0.002 | 0.126 ± 0.002 | 0.100 ± 0.003 | 0.147 ± 0.002 | -0.654 ± 0.015 | 0.244 ± 0.004 |
| ProteoGAN (100 labels) | 0.036 | 0.024 | 0.585 | 0.736 | -0.026 | 0.019 |
| ProteoGAN (200 labels) | 0.162 | 0.112 | 0.374 | 0.524 | 0.104 | 0.051 |

ProteoGAN clearly outperforms the HMM, n-gram model and CVAE on all metrics and embeddings. The same applies to the OpL versions which are trained once per label. ProteoGAN also outperforms the state-of-the-art ProGen model. MMD values are similar and ProGen would likely scale better than ProteoGAN, however MRR shows a clear advantage of ProteoGAN on conditional generation. We hypothesize that this is due to the stronger inductive bias of our conditioning mechanism.

The different embeddings (Tables 2,S6,S7) largely agree with each other, although it should be noted that the n-gram model almost directly optimizes the MMD-based metrics and is hence overestimated there, ProFET and UniRep provide a better picture in this case. The ESM embedding results (Table S8) were inconclusive as it indicated failure of all models. It also showed general instability with respect to model ranking depending on the choice of kernel parameters and homology levels (compare supplemental sections 11, 10, 15).

### 4.4 Ablation models: Hierarchical information dramatically improves conditioning

We also investigated several ablation models to demonstrate the advantages of conditioning on hierarchical labels. To begin with, the predictor-guided model had a very low conditional performance (MRR=0.114) and hence the original model exceeded it by a large margin (MRR=0.554). This shows that general function predictors for proteins are not (yet) suited to guide generative models at evaluation time. Continuous guidance during training might improve this result, but is time-wise prohibitive. The better MMD scores of the predictor model is likely due to the missing burden of the conditioning mechanism. We also observed this trade-off between MMD and MRR in the hyperparameter optimization.

Similarly, the non-hierarchical model (MMD=0.337, MRR=0.306) is clearly outperformed by the original ProteoGAN. The hierarchical information drastically helps model performance, presumably because the label structure can be transferred to the underlying sequence data structure, and because such a model does not need to learn each label marginal distribution independently.

The OpL-GAN separates the conditional learning problem into several sub-problems, and was in fact slightly better than ProteoGAN. Yet, ProteoGAN could achieve the result of 50 independent models by training a single conditional model with a minor trade-off in performance. Besides the lower training effort of ProteoGAN, the conditioning mechanism has the advantage to allow for functional OOD generation, as discussed below.

Figure 2 breaks down the conditional performance of ProteoGAN with respect to the individual labels. 27 of the 50 labels were on average ranked first or second and hence could very well be targeted. 33 of the 50 labels were ranked at least third. Certain branches of the ontology were more difficult to model (details available in Figure S9).

**Fig. 2:**
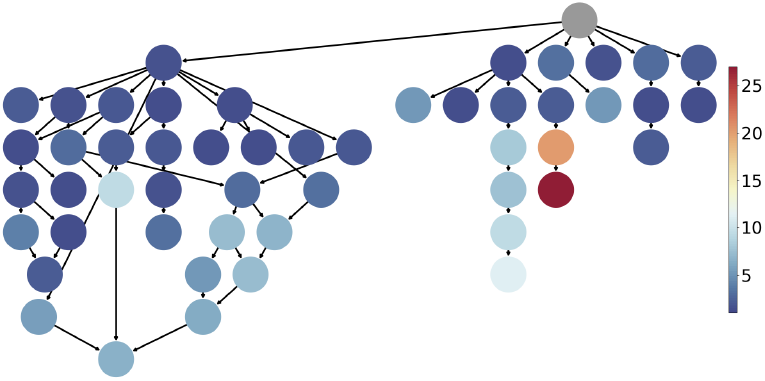
Mean rank of each individual label in Spectrum MRR over five data splits. The structure represents the relations of the 50 labels of interest in the GO DAG. Nodes are colored by how well the model can target them. Dark blue indicates that the model can target the function well, the worst targeted label (red) is “kinase activity”.

We asked how well our model could scale beyond 50 labels and trained it without any further tuning on 100 and 200 labels. While performance even gets better with 100 labels, once more showing that label information is advantageous, it starts to drop at 200 labels. Any scaling beyond this point will require hyperparameter tuning and likely an increase in parameter capacity to model the additional labels. Also, the amount of available training samples per class drops rapidly with increasing specificity of the labels. However the functional diversity we consider here would already enable many applications in *de novo* protein design.

### 4.5 Applicability: ProteoGAN can support protein screenings with a larger sequence space

It is difficult to prove biological validity without wet lab validation, and we do not claim to do so here. We acknowledge that MMD values still show significant difference to the positive control, and that corresponding p-values (Table S10) were inconclusive in this regard. Hence it is likely that generated sequences are not immediately usable out of the box, but need some experimental tuning as in directed evolution. Here we see the main application of ProteoGAN at this time: The extension of protein screenings with candidates that are further away from the known sequence space than previously possible, yet more likely to be functional than comparably novel candidates of other methods. To this end we compare ProteoGAN to random mutagenesis, the traditional method to produce candidates for such screenings, by gradually introducing random mutations into a set of natural sequences, simulating random mutagenesis with different mutation rates. We then compared MMD values between the mutated sets and generated sets from ProteoGAN (Table S23).

We first observed that generated sequences had an average maximum percent identity of 56% ± 2%, indicating that we are not simply reproducing training examples and sequences are novel. We refer in passing to Repecka *et al*. (2021) who had great success in validating GAN-generated proteins in vitro with a similar percent identity. Random mutagenesis with 90% sequence identity achieved the same MMD values as ProteoGAN, indicating that ProteoGAN is able to introduce four to five times more changes into the sequence at the same distributional shift. We conclude that ProteoGAN enables the exploration of a broader sequence space than random mutagenesis alone.

We further investigated how realistic the judgement of conditional performance by MRR is, by replacing the labels of generated sequences with the labels of their closest natural homolog (smallest edit distance). Interestingly, MRR remained high (MRR=0.379), despite low sequence similarity of the homologs. This shows that the sequences generated by ProteoGAN closely match the functional labels they were conditioned on, also when assessed by sequence similarity to known proteins.

### 4.6 Outlook: Conditioning may enable the design of novel protein functions

As an interesting outlook we provide first evaluations with respect to OOD generation. Models that condition on multiple labels generally aim to model the joint distribution of proteins given the labels, that is, proteins performing all indicated functions. We thus hypothesize that the conditioning mechanism may be used to combine previously unrelated functional labels into one protein, which would enable the design of completely novel kinds of proteins with previously unseen functionality. We stress that this objective is not explicitly build into the conditioning mechanism and thus it is not suited for the optimization of conflicting properties. However, combination of orthogonal properties might be permissive. While also here, biological implementation is inevitable to proof this concept, we can report that ProteoGAN and CVAE showed promising Top-X accuracies on five held out label combinations (Figures 3, S18). Further development of this concept will provide new tools for biotechnology.

**Fig. 3:**
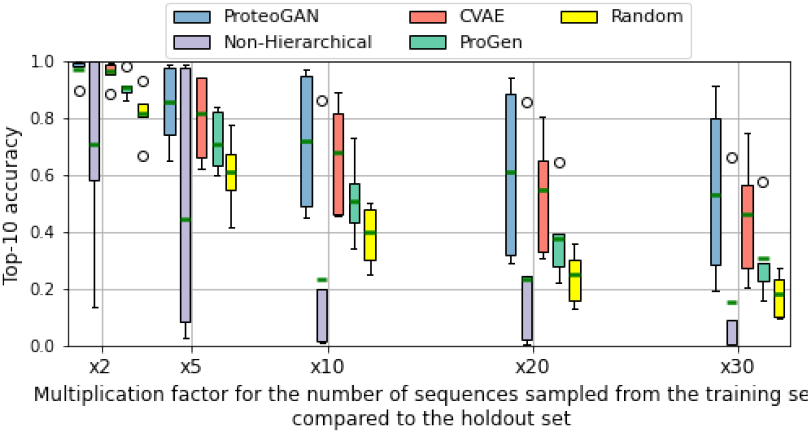
Top-10 accuracy (in %) with the Spectrum embedding for OOD-capable models. The boxplots cover the 5 OOD sets, A to E, the green line represents the average. We add a random baseline for comparison, where generated sequences are sampled uniformly at random from the training set. Complementary results for the other embeddings can be found in Figure S18.

### 4.7 Conclusion

We provide and evaluate broadly applicable metrics for assessing distribution similarity, conditional consistency and diversity in generative protein design. Due to their computational efficiency they can be used to compare and develop generative models at scale. With these metrics we hope to simplify the process of developing generative models in the domain of protein sequences. We present ProteoGAN, a GAN conditioned on hierarchical labels from the Gene Ontology, which outperforms classic and state-of-the-art models for (conditional) protein sequence generation. We envision that ProteoGAN may be used to exploit promising regions of the protein sequence space that are inaccessible by experimental random mutagenesis. It is universally applicable in various contexts that require different protein functions, and is even able to provide sequence candidates for never seen proteins. Extensions to this framework could incorporate other conditional information, such as structure motifs, binding partners, or other types of ontologies. Further development of such models may make proteins available as universal molecular machines that can be purely computationally designed *ad hoc* for any given biotechnological application.

## Acknowledgements

We thank Karsten Borgwardt for insightful discussions.

## Notes

### Competing Interest Statement

The authors have declared no competing interest.

