## Supplemental Information for "Conditional Generative Modeling for De Novo Protein Design with Hierarchical Functions"

#### Contents


1 Definition of $F_{\max}$-score 2

2 Conditioning mechanism 3

3 Data 5

4 Baselines 7

5 Hyperparameter search 8

6 Hyperparameter analysis 10

7 Losses, Duality Gap and real-time evaluation of ProteoGAN 15

8 Control for overfitting 16

9 Breakdown of conditional performance per label 17

10 Model evaluation with ProFET, UniRep and ESM. 18

11 Robustness and selection of embedding 19

12 P-values corresponding to MMD tests 22

13 Kolmogorov-Smirnov statistics 23

14 PCA of generated vs real sequences 25

15 Model evaluation with homology-controlled test sets 26

16 Comparison with random mutagenesis 30

17 OOD evaluation 31

18 Multiple Sequence Alignments of some generated sequences 32

19 Structure predictions of ProteoGAN generated proteins 33

#### 1 Definition of $F_{\max}$-score

The CAFA challenge [18] uses the $F_{\max}$-score to compare the performance of protein function predictors:

$$\begin{aligned}
pr(\tau) &= \frac{1}{m(\tau)} \sum_{i=1}^{m(\tau)} \frac{\sum_f \mathbb{1}(f \in P_i(\tau) \wedge f \in T_i)}{\sum_f \mathbb{1}(f \in P_i(\tau))}, \\
rc(\tau) &= \frac{1}{n} \sum_{i=1}^{n} \frac{\sum_f \mathbb{1}(f \in P_i(\tau) \wedge f \in T_i)}{\sum_f \mathbb{1}(f \in T_i)}, \\
F_{max} &= \max_{\tau} \left\{ \frac{2 \cdot pr(\tau) \cdot rc(\tau)}{pr(\tau) + rc(\tau)} \right\},
\end{aligned} \tag{1}$$

where $f$ is a GO term, $n$ is the number of sequences, $m(\tau)$ is the number of sequences with a score greater than or equal to $\tau$, $P_i(\tau)$ is the set of predicted terms with a score greater than or equal to $\tau$ for a protein sequence $i$, $T_i$ denotes the corresponding ground truth set of terms for that sequence, and $\mathbb{1}(\cdot)$ an indicator function.

#### 2 Conditioning mechanism

In this section, we first detail how we adapted the Wasserstein loss to the conditional setting, then we describe state-of-the-art conditional GAN objective functions and variants used in this project.

**Loss function of conditional GANs:** Our models are trained with the Wasserstein objective with gradient penalty from [7]. As a reminder, the WGAN-GP losses can be written as follows:

$$\begin{aligned}\mathcal{L}_D &= \mathbb{E}_{q(\boldsymbol{x})}[D(\boldsymbol{x})] - \mathbb{E}_{p(\boldsymbol{x})}[D(\boldsymbol{x})] \\ &\quad + \lambda\,\mathbb{E}_{m(\hat{\boldsymbol{x}})}[(\|\nabla_{\hat{\boldsymbol{x}}} D(\hat{\boldsymbol{x}})\|_2 - 1)^2] \\ \mathcal{L}_G &= -\,\mathbb{E}_{q(\boldsymbol{x})}[D(\boldsymbol{x})]\end{aligned} \tag{2}$$

where $\boldsymbol{x} \sim p(\boldsymbol{x})$ is the data distribution and $\boldsymbol{x} \sim q(\boldsymbol{x})$ is the generator model distribution, $\hat{\boldsymbol{x}}$ is an interpolated sample between a real sequence and a generated one, $m$ is the distribution of interpolated samples, $D$ is the discriminator (or critic), $\mathcal{L}_D$ the loss of the discriminator and $\mathcal{L}_G$ the loss of the generator. The term $\mathbb{E}_{m(\hat{\boldsymbol{x}})}[(\|\nabla_{\hat{\boldsymbol{x}}} D(\hat{\boldsymbol{x}})\|_2 - 1)^2]$ ensures that the discriminator is Lipschitz continuous.

To be able to use the Wasserstein objective with gradient penalty in the projection discriminator [13] (see below), we had to adapt the objective formula to include the label information. Let $D$ be the discriminator and $G$ the generator. Let $(\boldsymbol{x}, \boldsymbol{y}) \sim p$ be a sample from the dataset, where $\boldsymbol{x}$ is a one-hot encoded sequence, and $\boldsymbol{y}$ an encoding of choice of the categorical label. Let $q$ be the generator model distribution, such that $\boldsymbol{y} \sim q(\boldsymbol{y})$ is defined by the user and, in practice, follows the distribution of the labels in the dataset, and $\boldsymbol{x} \sim q(\boldsymbol{x}|\boldsymbol{y})$ is learned. Let $\hat{\boldsymbol{x}}$ be a linear interpolation between a real sequence and a generated one, with interpolation parameter chosen uniformly at random between 0 and 1. We call $\hat{\boldsymbol{x}} \sim m(\hat{\boldsymbol{x}}|\boldsymbol{y})$ the conditional distribution of interpolated sequences given a label encoding $\boldsymbol{y}$. Let $\lambda$ be a weighing factor introduced in [7]. The discriminator and generator losses can be written as follows:

$$\begin{aligned}\mathcal{L}_D &= \mathbb{E}_{q(\boldsymbol{y})}[\mathbb{E}_{q(\boldsymbol{x}|\boldsymbol{y})}[D(\boldsymbol{x}, \boldsymbol{y})]] - \mathbb{E}_{p(\boldsymbol{y})}[\mathbb{E}_{p(\boldsymbol{x}|\boldsymbol{y})}[D(\boldsymbol{x}, \boldsymbol{y})]] \\ &\quad + \lambda\,\mathbb{E}_{p(\boldsymbol{y})}[\mathbb{E}_{m(\hat{\boldsymbol{x}}|\boldsymbol{y})}[(\|\nabla_{\hat{\boldsymbol{x}}} D(\hat{\boldsymbol{x}}, \boldsymbol{y})\|_2 - 1)^2]], \\ \mathcal{L}_G &= -\,\mathbb{E}_{q(\boldsymbol{y})}[\mathbb{E}_{q(\boldsymbol{x}|\boldsymbol{y})}[D(\boldsymbol{x}, \boldsymbol{y})]].\end{aligned} \tag{3}$$

This formulation ensures that the Lipschitz constraints imposed on the discriminator in the unconditional WGAN-GP objective holds for each class.

**Case of the cGAN with projection discriminator [13]:** In the conditional GAN with projection discriminator model, the discriminator is decomposed into a sum of two terms, one being the inner product between a label embedding and an intermediate transformation of the input, and the second term being solely depending on the input sequence $\boldsymbol{x}$. The new expression of the projection discriminator can be derived by assuming that the label is categorical and that both the log-likelihoods of the data and target distribution can be written as log linear models. Let $\boldsymbol{y} \rightarrow \boldsymbol{v}(\boldsymbol{y})$ be a linear projection of the label encoding. Let $\boldsymbol{\phi}_\theta$ be an embedding function applied to the input $\boldsymbol{x}$ and $\psi_\gamma$ a scalar function applied to the embedding function $\boldsymbol{\phi}_\theta$. Let $\mathcal{A}$ be an activation function of choice. The projection discriminator in [13] can therefore be written as:

$$D(\boldsymbol{x}, \boldsymbol{y}) = \mathcal{A}(\boldsymbol{v}(\boldsymbol{y})^T \boldsymbol{\phi}_\theta(\boldsymbol{x}) + \psi_\gamma(\boldsymbol{\phi}_\theta(\boldsymbol{x}))) \tag{4}$$

The label information is therefore introduced via an inner-product. This formulation leads to a more stable algorithm compared to a simple concatenation of the label with the input [12], potentially thanks to the introduction of a form of regularization on the discriminator.

In this project, we tested the possibility to include several projections in the discriminator. In addition to the previous notations introduced in this section, let us assume that we have now $k$ projections. Let $\{g_i\}_{i=1}^k$ be $k$ neural networks, which can be decomposed in $n_i$ layers $g_i = l_{n_i}^i \circ \boldsymbol{l}_{n_i-1}^i \circ \cdots \boldsymbol{l}_2^i \circ \boldsymbol{l}_1^i$. Let $\{p_i\}_{i=1}^k$ be the layer number at which the inner product with the output of the linear projection $\{\boldsymbol{v}_i\}_{i=1}^k$ occurs in each neural network. The projections obey a tree-like branching structure, where all layers $p \leq p_i$ of the neural network $i$ are shared with the neural networks $j$ for which $p_i < p_j$ and the branching of a different projection is always done at a different layer number. The discriminator with multiple projections can then be written as:

$$D(\boldsymbol{x}, \boldsymbol{y}) = \mathcal{A}(\sum_{i=1}^{k} (\boldsymbol{v}_i(\boldsymbol{y})^T \boldsymbol{l}_{p_i}^i \circ \cdots \boldsymbol{l}_1^i(\boldsymbol{x}) + g_i(\boldsymbol{x}))) \tag{5}$$

In practice we allow for up to four projections. Our BOHB hyperparameter searches did not show evidence of the superiority of projection mechanisms for conditioning purposes when they are the unique type of conditional mechanism in the network. However, the projection models were able to generate sequences similar to naturally occurring ones (low MMD), and the resulting model of the optimization had two projections.

**Case of the cGAN model with auxiliary classifier [15]:** As opposed to cGANs with projection discriminator, cGANs with auxiliary classifier add a term to the generator and discriminator losses to incorporate the log-likelihood of the correct labels (compare Equation 3). In addition to notations introduced for Equation 3, let $C_D$ be the auxiliary classifier, $ce$ the cross entropy and $\gamma$ a weighting factor. For each label $\boldsymbol{y}$, the loss function of cGANs with auxiliary classifiers can be written as:

$$\begin{aligned}\mathcal{L}_D &= \mathbb{E}_{q(\boldsymbol{y})}[\mathbb{E}_{q(\boldsymbol{x}|\boldsymbol{y})}[D(\boldsymbol{x},\boldsymbol{y})]] - \mathbb{E}_{p(\boldsymbol{y})}[\mathbb{E}_{p(\boldsymbol{x}|\boldsymbol{y})}[D(\boldsymbol{x},\boldsymbol{y})]] \\ &\quad + \lambda\,\mathbb{E}_{p(\boldsymbol{y})}[\mathbb{E}_{m(\hat{\boldsymbol{x}}|\boldsymbol{y})}[(\|\nabla_{\hat{\boldsymbol{x}}} D(\hat{\boldsymbol{x}},\boldsymbol{y})\|_2 - 1)^2]] \\ &\quad + \gamma\,\mathbb{E}_{p(\boldsymbol{x}|\boldsymbol{y})}[ce(C_D(\boldsymbol{x}),\boldsymbol{y})] + \gamma\,\mathbb{E}_{q(\boldsymbol{x}|\boldsymbol{y})}[ce(C_D(\boldsymbol{x}),\boldsymbol{y})], \\ \mathcal{L}_G &= -\,\mathbb{E}_{q(\boldsymbol{y})}[\mathbb{E}_{q(\boldsymbol{x}|\boldsymbol{y})}[D(\boldsymbol{x},\boldsymbol{y})]] + \gamma\,\mathbb{E}_{p(\boldsymbol{x}|\boldsymbol{y})}[ce(C_D(\boldsymbol{x}),\boldsymbol{y})] + \gamma\,\mathbb{E}_{q(\boldsymbol{x}|\boldsymbol{y})}[ce(C_D(\boldsymbol{x}),\boldsymbol{y})]\end{aligned} \tag{6}$$

$C_D$ typically shares weights with $D$ and is trained when minimising $\mathcal{L}_D$ but is fixed when minimising $\mathcal{L}_G$.

In our work, we compare both types of conditional GANs (GAN equipped with auxiliary classifier or with multiple projections at several layers (see Equation 5)) to a third proposed model that combines both mechanisms. It is important to note that in this case the label information introduced in the projection may not be shared with the auxiliary classifier. The fANOVA analysis performed on the second BOHB optimization results (Figure S4) shows that the combination of both mechanisms helps to obtain a better performing conditioning mechanism, as measured by MRR.

### 3 Data

**Sequence data** was obtained from the UniProt Knowledgebase on May 23rd, 2020 via the query `(existence:"evidence at transcript level" OR existence:"evidence at protein level") goa:(evidence:manual) go:0003674`. For this proof-of-concept work we selected highly confident annotations and sequences, future work may profit from e.g. pretraining with a larger sequence pool.

**Functional labels** were collected from the same database. The gene ontology (GO) resource is composed of three branches, molecular function, cellular component and biological process. We focus on the molecular function ontology which contains thousands of terms ranging from description like *binding* (GO:0005488) to very specific terms such as *microtubule-severing ATPase activity* (GO:0008568). Each protein is annotated with a set of GO labels describing the molecular function of a protein in modular way. The ontology is structured as a directed acyclic graph with a single root. Further, labels have a hierarchical relationship, i.e. protein with a given functional label inherits automatically the labels of its parents in the DAG (*is-a* relationship). The molecular function ontology resource currently contains more than ten thousand labels, many of which have a highly specific meaning and only few representatives. We therefore restrict the number of labels to the 50 largest classes. We argue fifty labels is sufficient for a proof-of-principle and would even enable the design of experimental assays for validation. Figure S1 illustrates the selected subset of labels and their relationships. The ontology itself (the DAG) was downloaded at the same time from a separate source: `http://purl.obolibrary.org/obo/go/go-basic.obo`

**Train, validation and test splits** were created to preferably represent all labels uniformly in the test and evaluation sets. This is complicated by the multi-label structure of the data and we hence resort to a heuristic approach. To create the sets, we randomly sample sequences until there is at least 1.300 (300 in the optimization) sequences per class, where at each step we only sample from sequences that have at least one underrepresented label. The selection of hyperparameters by the BOHB hyperparameter optimizations for ProteoGAN and by the hyperparameter searches for the baselines are done on the validation set, while the results presented in the main text were acquired on the test set. For sequence sample generation the model was conditioned on the label combinations of the evaluation/test set and the resulting sequences then compared with the respective set by MMD and MRR.

**OOD holdout sets** were manually selected based on their usefulness in real-world application and the amount of samples they comprise (Table S2).

Table S1: Physicochemical properties and their accession numbers in the AAIndex.

| Property name | Accession number |
|---|---|
| Molecular Weight | FASG760101 |
| Hydrophobicity | FASG890101 |
| Hydrophilicity | HOPT810101 |
| Polarity | RADA880108 |
| Amphiphilicity | MITS020101 |
| Positive Charge | FAUJ880111 |
| Negative Charge | FAUJ880112 |
| pK | JOND750102 |
| Isoelectric Point | ZIMJ680104 |
| Probability Helix | KANM800101 |
| Probability Sheet | KANM800102 |
| Average Accessible Surface Area | JANJ780101 |
| Buriability | ZHOH040103 |
| Linker Index | BAEK050101 |

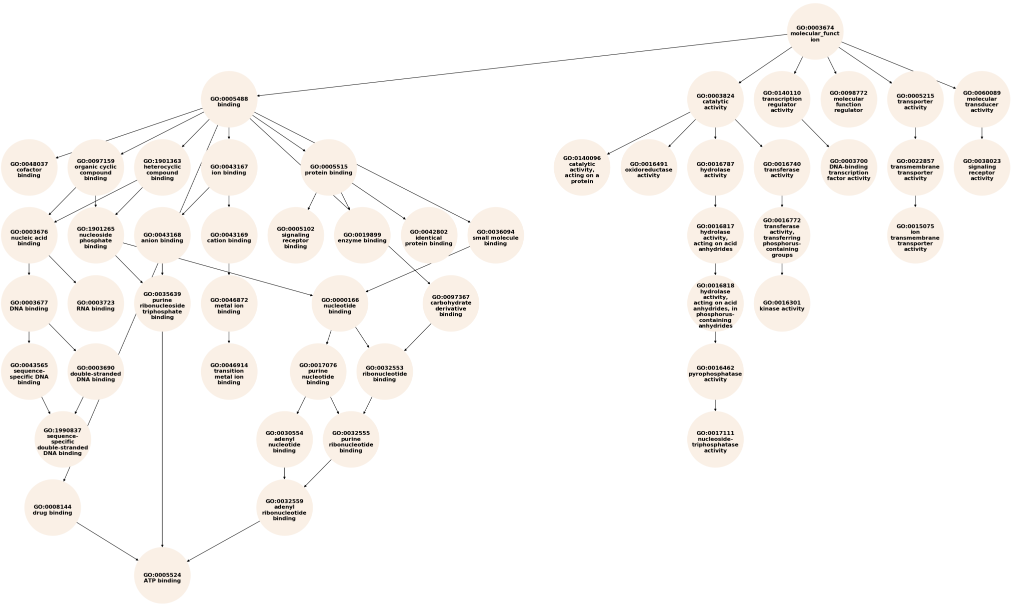


Figure S1: GO DAG of the 50 labels selected for this project.

Table S2: Label combinations held-out from the training dataset for OOD evaluation. Also given are the size of smallest class in $X_{\text{Train}}$ ($\text{Size}_{\text{Train}}$), the number of samples in $X_{\text{Holdout}}$ ($\text{Size}_{\text{Holdout}}$), and the average maximum percent identity of holdout sequences with the remaining dataset, with standard deviation.

| ID | Labels | $\text{Size}_{\text{Train}}$ | $\text{Size}_{\text{Holdout}}$ | Identity (%) |
|---|---|---|---|---|
| A | drug binding & transmembrane transporter activity | 10446 | 1678 | 62 ± 14 |
| B | catalytic activity & cofactor binding & anion binding | 5427 | 4027 | 78 ± 17 |
| C | small molecule binding & signaling receptor activity & catalytic activity, acting on a protein & kinase activity | 7676 | 1079 | 71 ± 17 |
| D | ion transmembrane transporter activity & cation binding | 7945 | 1109 | 62 ± 10 |
| E | DNA binding & RNA binding | 13207 | 1435 | 63 ± 13 |

### 4 Baselines

We implement several baselines to put the performance of our model into perspective. In this section, we would like to give additional details concerning the baselines that we gathered from the literature.

The HMM baselines were implemented based on HMMER [3]. For *OpC-HMM*, all sequences in the training dataset containing a specific *label combination* were aggregated, for each of the ca. 1.800 label combinations of the test set. For *OpL-HMM*, all sequences in the training dataset containing a specific *label* were aggregated, for each of the 50 labels. The resulting sequence sets were aligned with MAFFT (with parameters `--retree 1 --maxiterate 0 --ep 0.123`). Because of the time-intensive multiple sequence alignment the sequences sets were randomly sampled to have a maximum size of 5000 sequences. From the alignment, a profileHMM was built with HMMER which was then sampled to generate a sequence.

In the n-gram baseline also, sequences were selected according to label sets (OpC) and single labels (OpL). Here the full data was used. $n$ was set to 3. The sequence lengths were sampled from the sequence length distribution of the training data.

CVAE [5] uses a conditional VAE (CVAE) in order to generate either metalloproteins with desired metal binding sites or fold properties. In the case of fold properties, the authors introduce iterative sampling and guidance steps in the latent space. The decoder and encoder are both MLPs and the number of layers is chosen with hyperparameter search. We introduced a KL-balancing term to stabilize training. We performed a Bayesian Optimization hyperparameter search for which we tried $1,000$ combinations of hyperparameters. Notably, we allowed for an optimization of network architecture by optimizing over the layer numbers for both encoder and decoder, and by optimizing the number of units in the first layer of the encoder and the last layer of the decoder. The unit number then halved towards the latent space with each layer. The hyperparameters and their value ranges, as well as the final model configuration can be found in Table S3. We refer the reader to Greener et al. [5] for more information on the model.

ProGen's source code was collected from `https://github.com/lucidrains/progen`. We adjusted the model size in order to have reasonable and comparable runtimes to the other models. We trained a 4-layer model with 4 attention heads of dimension 66 in the heads and dimension 132 in the linear layers with window size 264 for 100 epochs with batch size 36. The model was trained and evaluated on the same dataset as the others. The perplexity of ProGen on our test set was 11.77.

Table S3: CVAE of Greener et al. hyperparameters subject to BO optimization.

| Name | | Values | Final Value |
|---|---|---|---|
| Learning rate | | [1e-5,1e-2] | 7.8e-4 |
| Pretrain start | | [1,5000] | 2598 |
| Pretrain end | | [1,5000] | 1251 |
| Latent dimension | | [10,1000]$^\dagger$ | 761 |
| KL balancing | $\beta$ | [1e-3,100]$^\dagger$ | 1.1e-3 |
| Encoder layer number | | [1,5] | 3 |
| Decoder layer number | | [1,5] | 1 |
| Log2(Encoder first layer units) | | [4,10] | 7 |
| Log2(Decoder last layer units) | | [4,10] | 9 |

$^\dagger$ Values were sampled on a logarithmic scale.

### 5 Hyperparameter search

For ProteoGAN we conducted hyperparameter searches with the Bayesian Optimization and HyperBand (BOHB) algorithm. The Hyperband [11] algorithm uses successive halving [9] to evaluate a number of models on a given budget of resources. The better half of the models are then evaluated on twice the budget, et cetera. Hyperband is an independent optimization algorithm that has been combined with Bayesian optimization to form Bayesian optimization and Hyperband (BOHB) [4], the optimization strategy used in this project.

We conducted two BOHB optimizations. For both we evaluated 1,000 models. All networks were trained with the Adam optimizer [10] with $\beta_1 = 0$ and $\beta_2 = 0.9$ (following [7]). The optimization consisted first of a broad search among 23 hyperparameters and second, of a smaller and more specific search, among 9 selected hyperparameters. For the first BOHB optimization, an optimization iteration was defined as two epochs which we found through pilot experiments was the minimum time to observe a viable trend in the metrics. The parameters $R$ and $\eta$ (in the notation of Li et al. [11]) were set to 9 and 3, respectively, which allowed for a maximum training time of 18 epochs (22.5K gradient updates). The optimization objective was to maximize the ratio of metrics MMD/MRR on the validation set (the metrics are introduced in the main document). During the optimization, BOHB selected the models based on evaluations at the end of a training period. For the second optimization, we reduced the number of hyperparameters to only 9. We selected values for the other hyperparameters based on the analysis of the hyperparameter importance of the first optimization (see paragraph below). The hyperparameters that showed either no importance or that were detrimental to training were removed. For this second optimization, the smaller network size allowed for 3 epochs per iteration, resulting in a maximum training time of 27 epochs (1.2K gradient updates). The list of hyperparameters of the two BOHB optimizations and their ranges is presented in Table S4. The parameters of the best models selected by the two BOHB optimizations are presented Table S5.

Table S4: Hyperparameters subject to BOHB optimization.

| Name | Symbol | Values |
|---|---|---|
| Use physicochemical properties | | Yes, No, only |
| Label embedding | | one-hot, node2vec, Poincaré |
| Conditioning mechanism | | projection, AC, both |
| AC weighting factor | $\gamma$ | [1, 1000]$^\dagger$ |
| Label smoothing factor | $\theta$ | [0, 0.5] |
| Latent noise dimension | $d_Z$ | [1, 1000]$^\dagger$ |
| Input noise standard deviation | $\sigma$ | [0, 1] |
| Generator learning rate | $\eta_G$ | [1e-5, 1e-2] |
| Generator learning rate 2 | $\eta_{G2}$ | [1e-5, 1e-2] |
| Discriminator learning rate | $\eta_D$ | [1e-5, 1e-2] |
| Discriminator learning rate 2 | $\eta_{D2}$ | [1e-5, 1e-2] |
| Training ratio | $n_{critic}$ | [1, 10] |
| Learning rate schedule | | constant, cosine, exponential |
| Schedule interval (in epochs) | $i$ | [1, 18] |
| Generator layer number | $n_G$ | [1, 10] |
| Discriminator layer number | $n_D$ | [1, 10] |
| Strides | $s$ | 1, 2, 4, 8 |
| Filter size | $f$ | [3, 12] |
| Generator skip | $h_G$ | [0, 10] |
| Discriminator skip | $h_D$ | [0, 10] |
| Number of projections | $n_P$ | [1, 5] |
| Output source layers | $o_S$ | [1, 3] |
| Output label layers | $o_L$ | [1, 3] |

$^\dagger$ Values were sampled on a logarithmic scale. AC = auxiliary classifier.

Table S5: Hyperparameters found in the first and second BOHB optimization. Values with an asterisk indicate the preset configurations in the second optimization.

| Name | First | Second |
|---|---|---|
| Use physicochemical properties | Yes | No |
| Label embedding | one-hot | one-hot |
| Conditioning mechanism | both | both |
| AC weighting factor | 178 | 135 |
| Label smoothing factor | 0.28 | -* |
| Latent noise dimension | 91 | 100* |
| Input noise standard deviation | 0.29 | -* |
| Generator learning rate | 2.0e-3 | 4.1e-4 |
| Generator learning rate 2 | - | -* |
| Discriminator learning rate | 8.5e-4 | 4.0e-4 |
| Discriminator learning rate 2 | - | -* |
| Training ratio | 1 | 1* |
| Learning rate schedule | constant | constant* |
| Schedule interval (in epochs) | - | -* |
| Generator layer number | 2 | 2* |
| Discriminator layer number | 3 | 2* |
| Strides | 4 | 8 |
| Filter size | 8 | 12 |
| Generator skip | - | -* |
| Discriminator skip | - | -* |
| Number of projections | 1 | 2 |
| Output source layers | - | 1* |
| Output label layers | 2 | 1* |

AC = auxiliary classifier.

### 6 Hyperparameter analysis

After the optimization we analyzed hyperparameter importance with the approach presented in [8]. A surrogate model (random forest) is trained on the parameter configurations and the respective evaluation scores. This enables a functional analysis of variance (fANOVA) which allows for a quantification of hyperparameter importance in terms of the variance they explain. It also provides marginal predictions for each hyperparameter which gives insights about their optimal value setting. For the random forest, we used $1,000$ trees with a maximum depth of 64, and repeat the estimation 100 times. We do so for all evaluated models of the first and second BOHB optimizations. The hyperparameter importances obtained from the first optimization (and resp. second optimization) are presented in Figure S5 (resp. Figure S6). The first fANOVA showed that parameters related to the discriminator (learning rate, number of layers) are most important for model performance,[1] and helped to select potentially important hyperparameters for the second analysis. Noticeably, the best model of the first optimization was already a well-performing model, but we chose to run a second optimization to better understand the role of key hyperparameters to gain insight in potential good practices when designing conditional generative adversarial networks. The second fANOVA clarified the importance of the remaining hyperparameters, such as use of physicochemical features and label embeddings among others (Figure S6).

We also show marginal predictions for hyperparameters of the first optimization in Figure S2, and for the second optimization in Figure S3 and Figure S4.

[1]Some other important factors were learning rate schedule-related parameters such as *Generator learning rate 2* or *schedule*. We realized that these were detrimental to model performance as the short duration of training in the optimization did not allow to estimate long term effects seen in the selected models that were trained for 100 epochs.

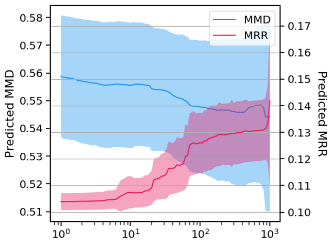


(a) Auxiliary Classifier weighing factor

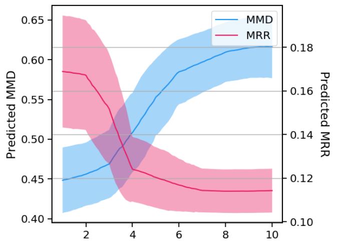


(b) Discriminator layer number

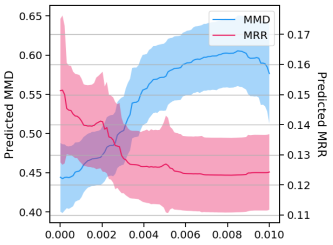


(c) Discriminator learning rate

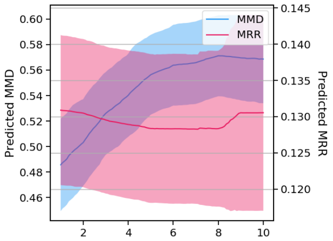


(d) Generator layer number

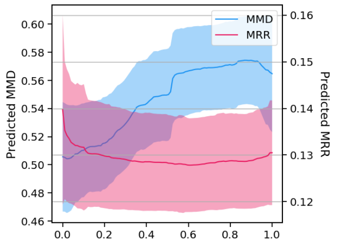


(e) Input noise

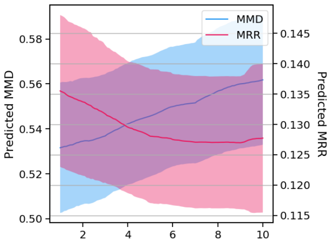


(f) Training ratio

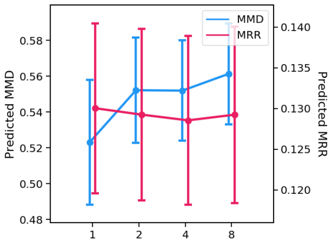


(g) Strides

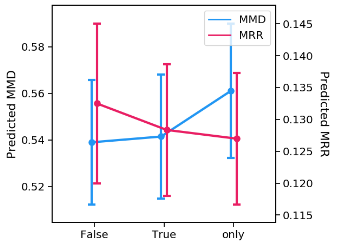


(h) Use physicochemical features

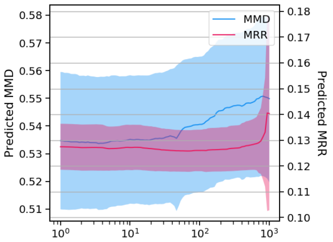


(i) Latent dimensionality

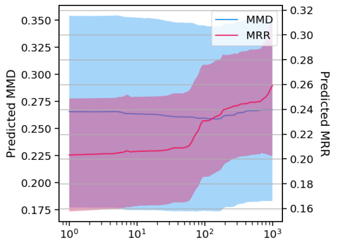


(j) Auxiliary Classifier weighing factor (Selected)

Figure S2: Marginal predictions of hyperparameters based on data in the first optimization. We show some selected predictions that allowed for interpretation, all others were inconclusive. If not otherwise noted, data comes from all trials in the optimization. Predictions were obtained training on MMD and MRR. Note that for MMD, lower is better.

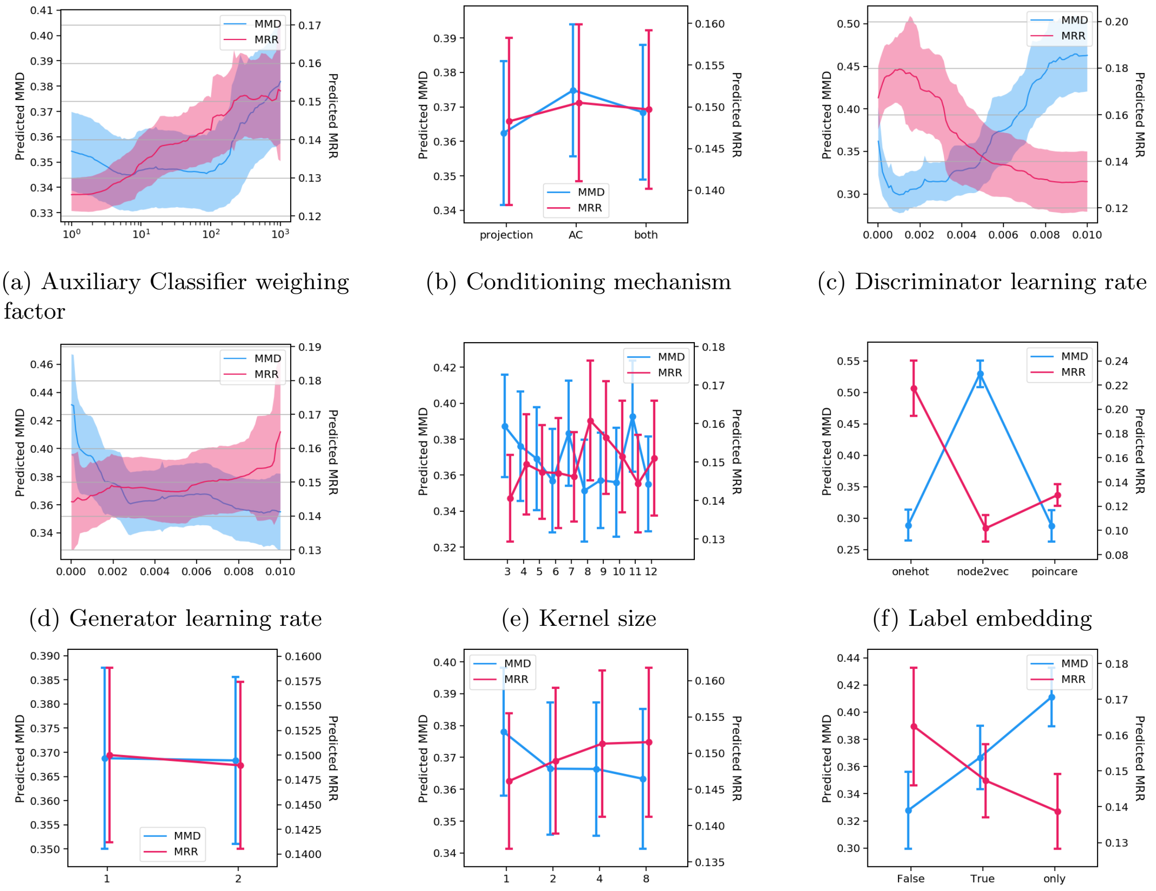


(a) Auxiliary Classifier weighing factor

(b) Conditioning mechanism

(c) Discriminator learning rate

(d) Generator learning rate

(e) Kernel size

(f) Label embedding

(g) Projections

(h) Strides

(i) Use physicochemical features

Figure S3: Marginal predictions of hyperparameters based on optimization data in the second optimization. Predictions were obtained training on MMD and MRR. Note that for MMD, lower is better.

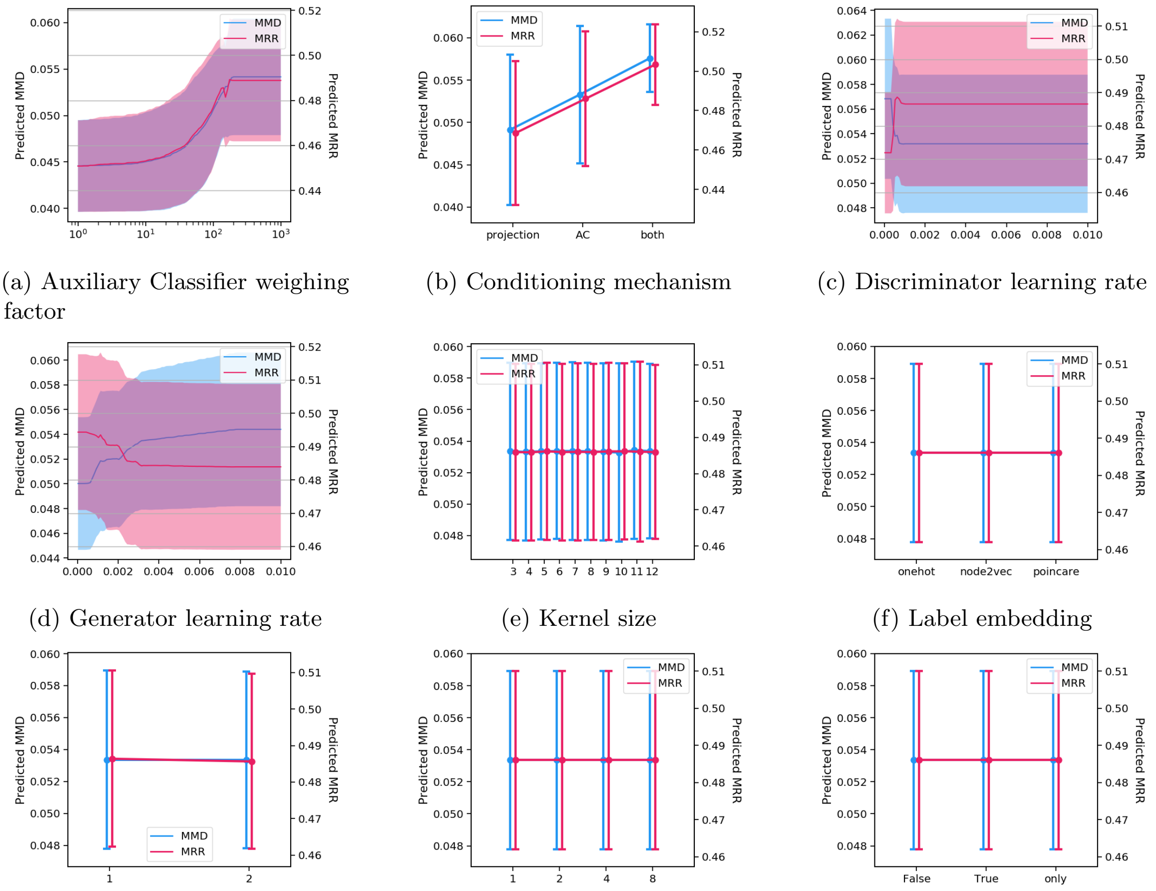


(a) Auxiliary Classifier weighing factor
(b) Conditioning mechanism
(c) Discriminator learning rate
(d) Generator learning rate
(e) Kernel size
(f) Label embedding
(g) Projections
(h) Strides
(i) Use physicochemical features

Figure S4: Marginal predictions of hyperparameters based on the data of the 27 best selected models in the second optimization. Predictions were obtained training on MMD and MRR. Note that for MMD, lower is better.

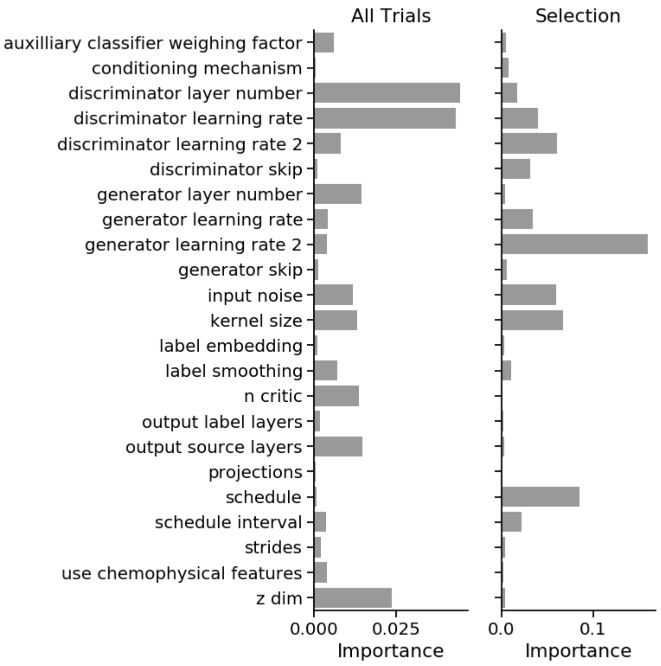


Figure S5: Hyperparameter importance for the first BOHB optimization. Shown are all hyperparameters subject to optimization for all models (left), and a manual selection of models that was trained for 100 epochs (right).

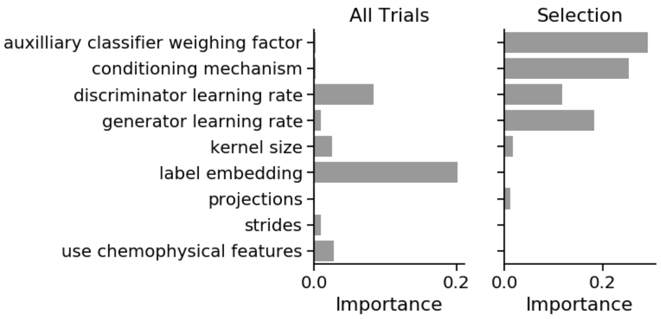


Figure S6: Hyperparameter importance for the second BOHB optimization. The bars show individual importance of each hyperparameter in terms of the variance they explain. We conducted the analysis for all trials of the optimization (left) and for the selected models that were trained for prolonged time (100 epochs) (right). The total variance explained by the main effects was 36% and 88%, respectively.

#### 7 Losses, Duality Gap and real-time evaluation of ProteoGAN

The loss function of the final model presented in the main document, combining projection and auxiliary classifier, is shown Figure S7. We monitored the duality gap (red), for which we split the training data into an adversary finding set and a test set of 1% of the train set each (these sets are required additionally to estimate the duality gap, compare Grnarova et al. [6]). The duality gap is well-behaved, with a fast convergence to 0, indicating that there is no mode collapse and suggesting that the samples are of reasonable quality. Also, the evaluations of MMD and MRR can be seen during training (evaluated twice per epoch) which provides valuable information for model selection and early stopping.

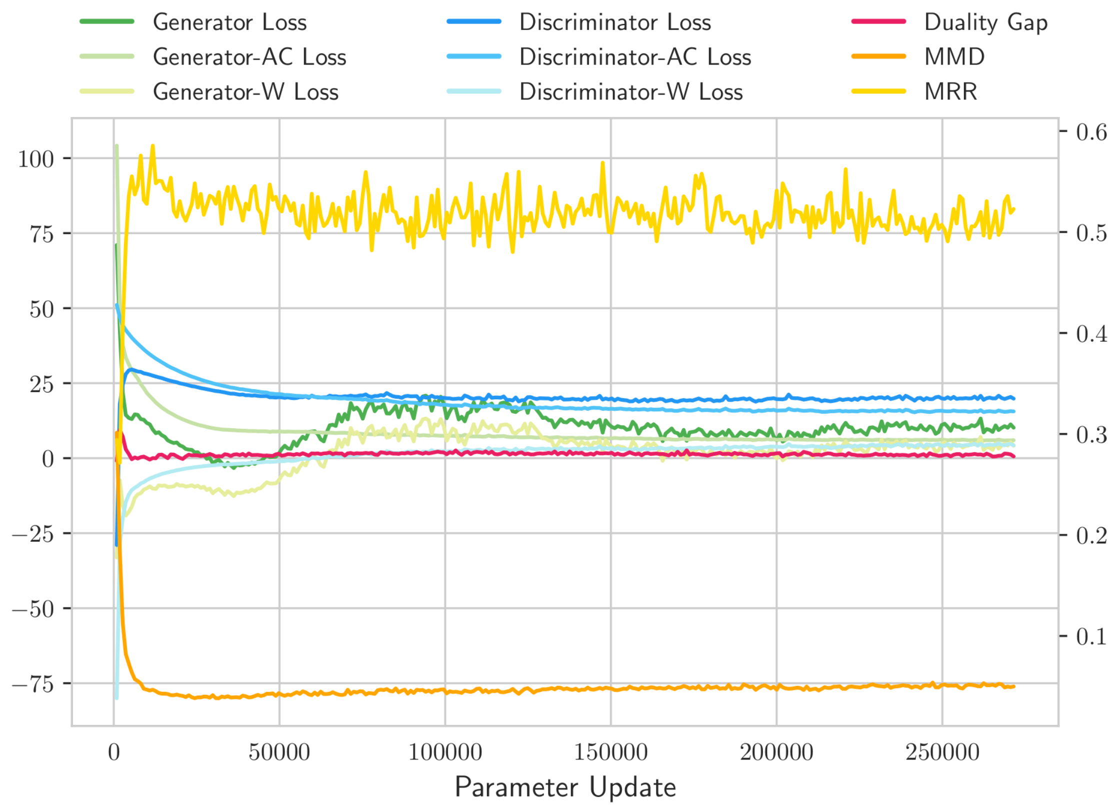


Figure S7: Losses and evaluations at training time. W = Wasserstein, AC = Auxiliary Classifier

### 8 Control for overfitting

To control that our model is not simply reproducing training examples we computed pairwise distances in feature space and verified that generated sequences were not closer to the training set than the test set (Figure S8).

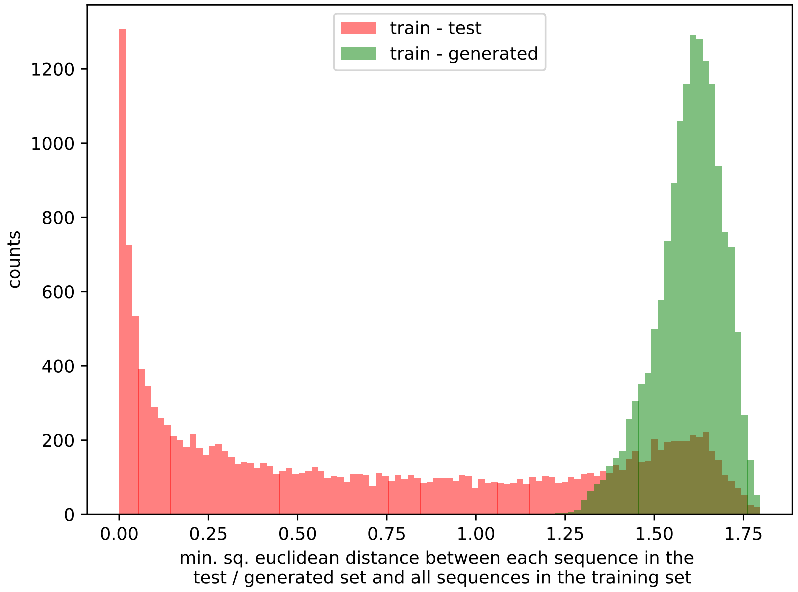


Figure S8: Distributions of pairwise distances in Spectrum kernel feature space between a test set and the training set (red) and a generated set of ProteoGAN and the training set (green). It can be seen that the generated sequences are not closer to the training set than the testset (which would indicate overfitting). Further the generated sequences are about as far, but not further, away from the training set than the testset.

### 9 Breakdown of conditional performance per label

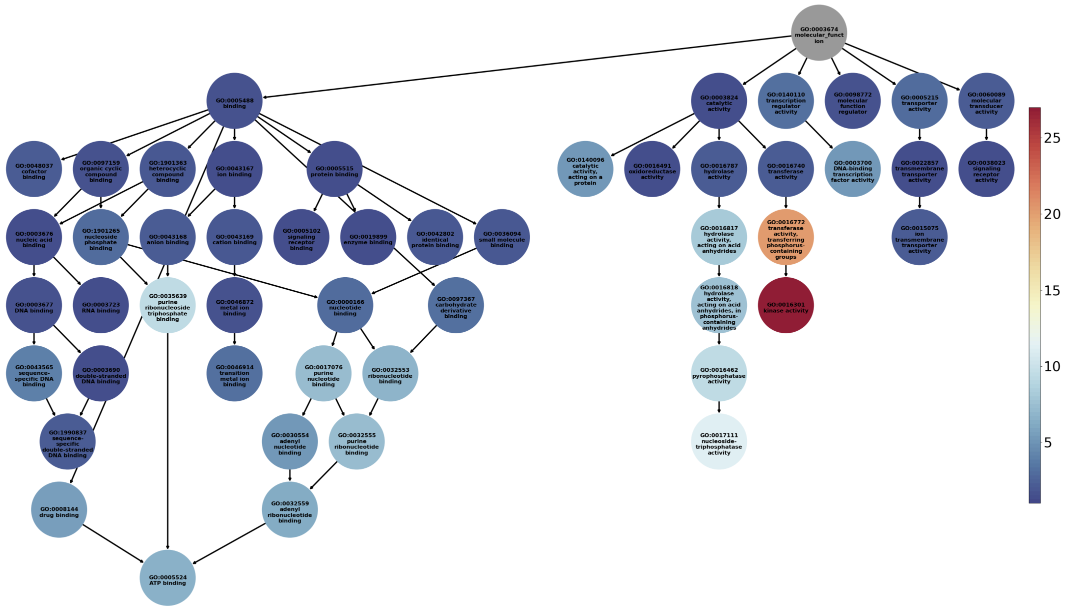


Figure S9: Mean rank of each individual label in Spectrum MRR over five data splits. The structure represents the relations of the 50 labels of interest in the GO DAG. Nodes are colored by how well the model can target them. Dark blue indicates that the model can target the function well.

### 10 Model evaluation with ProFET, UniRep and ESM.

Table S6: Evaluation of ProteoGAN and other models with proposed metrics, based on the ProFET embedding (compare results in the main text).

| Model | MMD↓ | gauss. MMD↓ | MRR↑ | $MRR_B$ ↑ | ΔEntropy | ΔDistance |
|---|---|---|---|---|---|---|
| Positive Control | 0.019 ± 0.005 | 0.011 ± 0.000 | 0.826 ± 0.021 | 0.928 ± 0.021 | -0.004 ± 0.006 | -0.010 ± 0.060 |
| Negative Control | 11.704 ± 0.005 | 1.006 ± 0.000 | 0.090 ± 0.000 | 0.119 ± 0.000 | 7.208 ± 0.010 | 6.373 ± 0.056 |
| ProteoGAN | 0.163 ± 0.014 | 0.038 ± 0.002 | 0.594 ± 0.050 | 0.766 ± 0.063 | -0.047 ± 0.025 | -0.328 ± 0.217 |
| Predictorguided | 0.091 ± 0.005 | 0.024 ± 0.001 | 0.131 ± 0.016 | 0.168 ± 0.018 | 0.032 ± 0.023 | 0.151 ± 0.147 |
| Non-Hierarchical | 1.144 ± 0.242 | 0.142 ± 0.016 | 0.288 ± 0.045 | 0.402 ± 0.075 | -0.215 ± 0.231 | -7.006 ± 4.127 |
| ProGen | 0.271 | 0.047 | 0.353 | 0.499 | 0.223 | 0.833 |
| CVAE | 1.086 ± 0.281 | 0.100 ± 0.006 | 0.286 ± 0.034 | 0.408 ± 0.039 | -0.145 ± 0.106 | -2.841 ± 2.097 |
| OpC-ngram | 0.548 ± 0.014 | 0.048 ± 0.001 | 0.259 ± 0.016 | 0.342 ± 0.022 | -0.005 ± 0.009 | -6.804 ± 0.264 |
| OpC-HMM | 0.689 ± 0.012 | 0.201 ± 0.002 | 0.170 ± 0.012 | 0.235 ± 0.013 | 1.135 ± 0.013 | 2.937 ± 0.057 |
| OpL-GAN | 0.161 | 0.043 | 0.644 | 0.807 | 0.196 | 0.981 |
| OpL-ngram | 0.580 ± 0.016 | 0.055 ± 0.001 | 0.267 ± 0.022 | 0.326 ± 0.021 | 0.022 ± 0.008 | -6.738 ± 0.327 |
| OpL-HMM | 0.798 ± 0.013 | 0.225 ± 0.003 | 0.176 ± 0.013 | 0.236 ± 0.014 | 1.330 ± 0.006 | 3.089 ± 0.038 |
| ProteoGAN (100 labels) | 0.124 | 0.037 | 0.537 | 0.746 | -0.041 | -0.547 |
| ProteoGAN (200 labels) | 0.245 | 0.079 | 0.297 | 0.412 | 0.314 | 0.856 |

Table S7: Evaluation of ProteoGAN and other models with proposed metrics, based on the UniRep embedding (compare results in the main text).

| Model | MMD↓ | gauss. MMD↓ | MRR↑ | $MRR_B$ ↑ | ΔEntropy | ΔDistance |
|---|---|---|---|---|---|---|
| Positive Control | 0.038 ± 0.005 | 0.011 ± 0.000 | 0.855 ± 0.040 | 0.946 ± 0.034 | -0.000 ± 0.004 | -0.050 ± 0.185 |
| Negative Control | 10.618 ± 0.002 | 1.000 ± 0.000 | 0.090 ± 0.000 | 0.119 ± 0.000 | 8.121 ± 0.007 | 27.981 ± 0.169 |
| ProteoGAN | 1.512 ± 0.019 | 0.061 ± 0.003 | 0.244 ± 0.013 | 0.300 ± 0.011 | 0.866 ± 0.027 | 12.611 ± 0.731 |
| Predictorguided | 1.591 ± 0.012 | 0.068 ± 0.002 | 0.097 ± 0.004 | 0.121 ± 0.005 | 0.964 ± 0.017 | 15.004 ± 0.263 |
| Non-Hierarchical | 2.408 ± 0.456 | 0.069 ± 0.009 | 0.170 ± 0.037 | 0.225 ± 0.034 | 0.689 ± 0.166 | 4.790 ± 3.039 |
| ProGen | 1.240 | 0.054 | 0.255 | 0.331 | 0.762 | 10.938 |
| CVAE | 1.808 ± 0.203 | 0.046 ± 0.010 | 0.213 ± 0.016 | 0.276 ± 0.017 | 0.632 ± 0.109 | 7.201 ± 3.752 |
| OpC-ngram | 2.062 ± 0.009 | 0.100 ± 0.001 | 0.147 ± 0.005 | 0.192 ± 0.014 | 0.943 ± 0.006 | 16.176 ± 0.161 |
| OpC-HMM | 2.083 ± 0.021 | 0.194 ± 0.002 | 0.116 ± 0.007 | 0.146 ± 0.011 | 1.529 ± 0.011 | 18.765 ± 0.182 |
| OpL-GAN | 1.721 | 0.095 | 0.224 | 0.278 | 1.108 | 16.853 |
| OpL-ngram | 2.154 ± 0.008 | 0.118 ± 0.001 | 0.159 ± 0.006 | 0.224 ± 0.008 | 1.025 ± 0.011 | 17.424 ± 0.185 |
| OpL-HMM | 2.263 ± 0.026 | 0.244 ± 0.006 | 0.120 ± 0.018 | 0.150 ± 0.017 | 1.749 ± 0.028 | 19.904 ± 0.428 |
| ProteoGAN (100 labels) | 1.446 | 0.055 | 0.225 | 0.294 | 0.820 | 10.707 |
| ProteoGAN (200 labels) | 2.059 | 0.149 | 0.126 | 0.177 | 1.393 | 21.001 |

Table S8: Evaluation of ProteoGAN and other models with proposed metrics, based on the ESM embedding (compare results in the main text).

| Model | MMD↓ | gauss. MMD↓ | MRR↑ | $MRR_B$ ↑ | ΔEntropy | ΔDistance |
|---|---|---|---|---|---|---|
| Positive Control | 0.045 ± 0.006 | 0.011 ± 0.000 | 0.868 ± 0.029 | 0.956 ± 0.010 | -0.001 ± 0.004 | 0.023 ± 0.202 |
| Negative Control | 10.303 ± 0.002 | 1.000 ± 0.000 | 0.090 ± 0.000 | 0.105 ± 0.001 | 8.808 ± 0.003 | 27.127 ± 0.074 |
| ProteoGAN | 5.751 ± 0.032 | 0.063 ± 0.004 | 0.095 ± 0.006 | 0.109 ± 0.006 | 0.729 ± 0.035 | 17.790 ± 0.488 |
| Predictorguided | 5.686 ± 0.016 | 0.083 ± 0.003 | 0.090 ± 0.001 | 0.104 ± 0.001 | 0.849 ± 0.014 | 19.212 ± 0.169 |
| Non-Hierarchical | 5.610 ± 0.198 | 0.058 ± 0.013 | 0.092 ± 0.002 | 0.106 ± 0.003 | 0.366 ± 0.149 | 8.590 ± 5.563 |
| ProGen | 5.463 | 0.098 | 0.091 | 0.104 | 0.925 | 19.563 |
| CVAE | 5.728 ± 0.097 | 0.025 ± 0.005 | 0.092 ± 0.002 | 0.106 ± 0.002 | 0.225 ± 0.137 | 6.509 ± 4.271 |
| OpC-ngram | 5.803 ± 0.019 | 0.071 ± 0.001 | 0.091 ± 0.000 | 0.104 ± 0.000 | 0.704 ± 0.002 | 16.852 ± 0.085 |
| OpC-HMM | 6.324 ± 0.018 | 0.170 ± 0.003 | 0.090 ± 0.000 | 0.104 ± 0.000 | 1.189 ± 0.008 | 22.038 ± 0.104 |
| OpL-GAN | 5.688 | 0.097 | 0.102 | 0.115 | 0.881 | 18.983 |
| OpL-ngram | 5.803 ± 0.020 | 0.082 ± 0.001 | 0.091 ± 0.000 | 0.104 ± 0.000 | 0.751 ± 0.005 | 17.453 ± 0.224 |
| OpL-HMM | 6.500 ± 0.019 | 0.202 ± 0.002 | 0.090 ± 0.000 | 0.104 ± 0.000 | 1.283 ± 0.004 | 22.912 ± 0.073 |
| ProteoGAN (100 labels) | 5.658 | 0.061 | 0.101 | 0.115 | 0.737 | 17.453 |
| ProteoGAN (200 labels) | 5.855 | 0.104 | 0.090 | 0.104 | 0.921 | 19.327 |

#### 11 Robustness and selection of embedding

There are various hyperparameters in MMD that need to be tuned, such as the embedding, the choice of kernel, and its parameters. Different settings can result in vastly different results, to the extent that models may be ranked differently. Selecting the right hyperparameters is challenging because the optimal set highly depends on the task at hand. O'Bray et al. [14] propose a method based on perturbations to select the best embedding, kernel and kernel parameters. Based on the desideratum that MMD shall detect small differences in the distributions of two samples behaving as a metric, they introduce increasing amounts of noise into a sample of real data. Ideally, MMD between a real sample and the perturbed samples should show a high correlation with the amount of perturbation added, and parameters can be selected on the highest correlation.

We performed such an analysis (Figure S10) with various, simple types of noises that we would expect in the context of protein generation (Table S9). We found that all embeddings were affected by the choice of kernel and parameters. However, the Spectrum embedding proved to be least affected. As opposed to the other embeddings, the linear kernel applied to the Spectrum embedding remained high across all types of perturbations (orange, Figure S10). As can be seen in Figure S11, the optimal parameters of the Gaussian kernel differ between types of perturbations. It is hence pertinent to choose the most robust embedding for the task of evaluation, the Spectrum kernel embedding. Note that the correlation is high in all cases, and the embedding is robust not because it is generally bad. We further show that euclidean distances of Spectrum embeddings correlate best with sequence alignment distances (Figure S12).

We think that the reason for the higher robustness of the Spectrum embedding is its simplicity. The other embeddings are complex, learned mappings of sequences that were trained on a very large set of natural sequences. As such, they (deliberately) overfit to natural sequences and behave badly on artificial sequences that do not follow the natural protein distribution (the problem of distribution shift). While we eventually aim to generate proteins that exactly follow this distribution, it is indispensable for a metric of generative models to be discriminative also in regimes that do not perfectly resemble the real distribution. The Spectrum embedding makes no assumption on the distribution of proteins and is hence better suited for the metrics we propose in this paper.

Table S9: Sequence- and sample-based perturbations. The severity of the perturbations were varied between 0% and 10% in 10 regularly spaced intervals. Severity is indicated as x.

| Perturbation | Description |
|---|---|
| Mutation | x% of residues in each sequence were mutated to a random amino acid |
| Insertion | x% of residues in each sequence were inserted |
| Deletion | x% of residues in each sequence were deleted |
| Random | x% of the sample was replaced with a random amino acid string |
| Chimera | x% of the sample was replaced with half-half chimeras of two proteins |
| Shorten | x% of the sample was shortened from the end by x% |

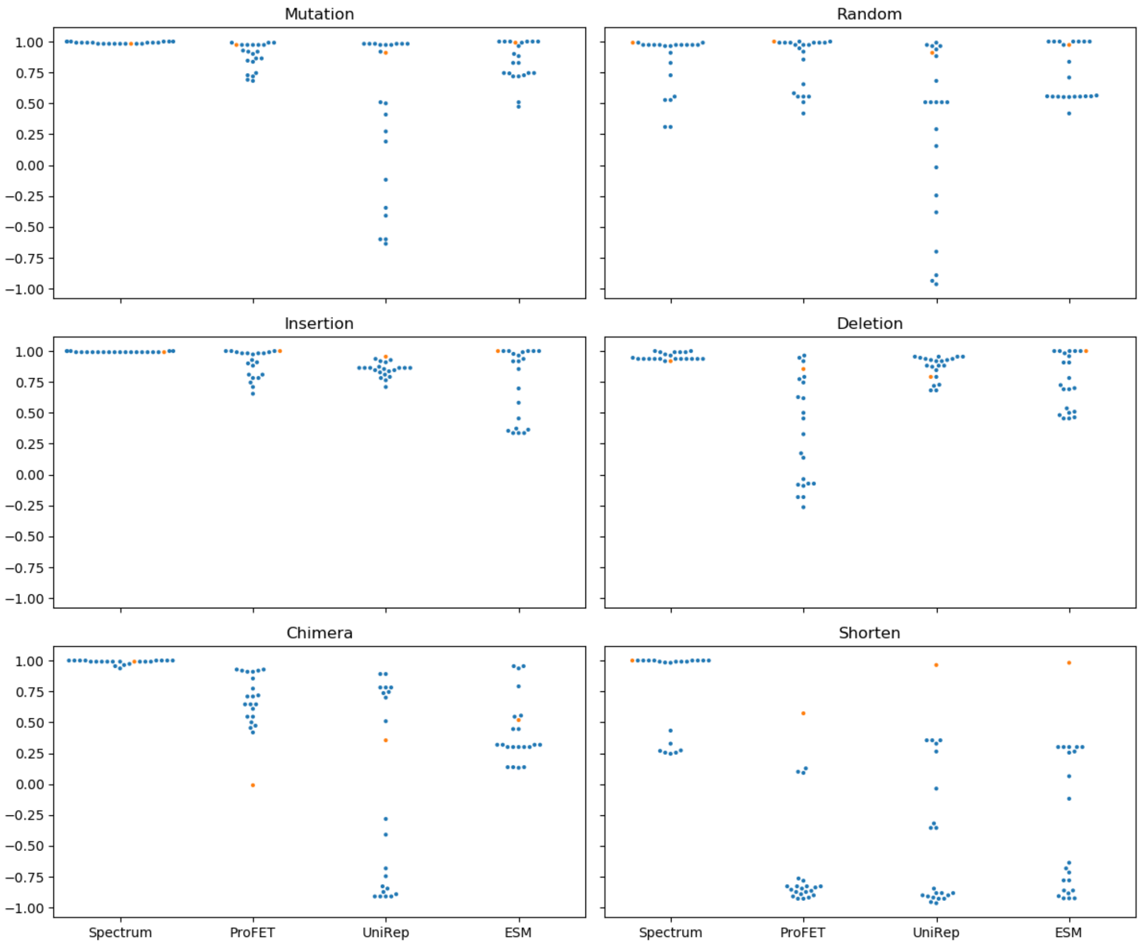


Figure S10: Spearman correlation of different perturbations with MMD based on different embeddings and kernel parameters over 1000 sequences. The linear kernel in orange, different Gaussian kernel parameters (gamma) in blue. Kernel parameters were varied between 0.1 and 10 times the median of embedding distances in the unperturbed set, in logarithmic intervals.

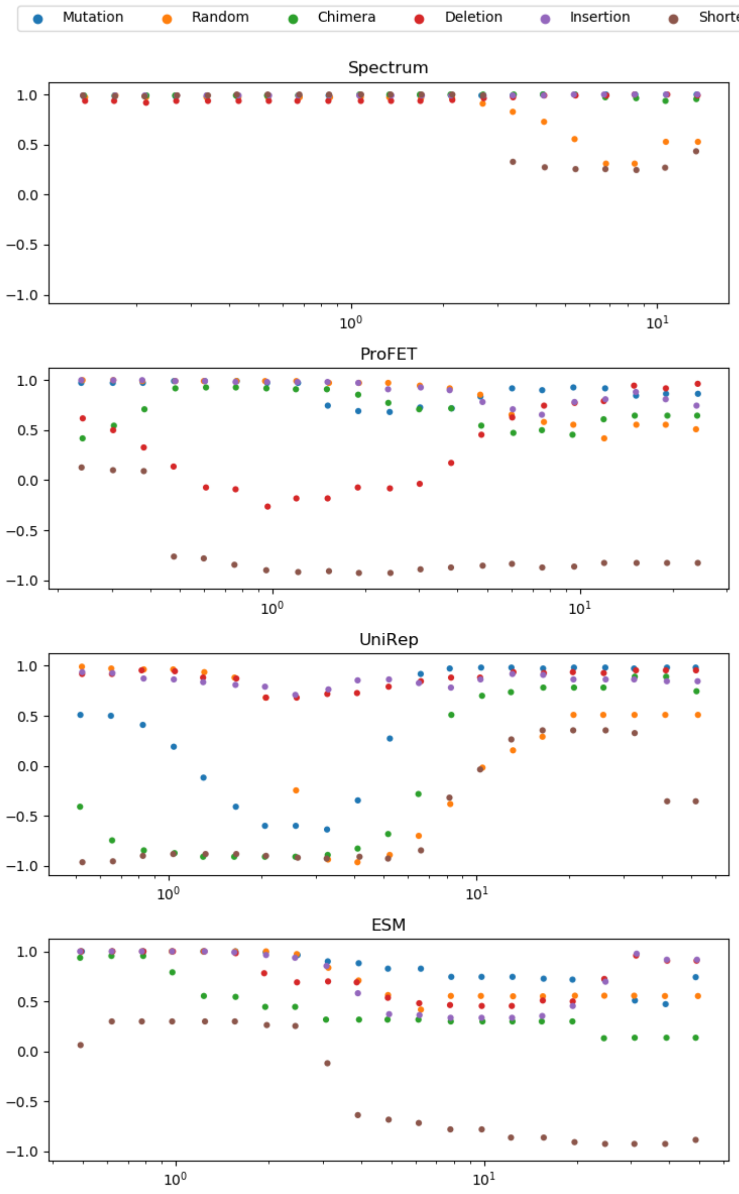


Figure S11: Spearman correlation of different perturbations with MMD based on different embeddings and kernel parameters over 1000 sequences. Kernel parameters (gamma, on the x-axis) were varied between 0.1 and 10 times the median of embedding distances in the unperturbed set, in logarithmic intervals.

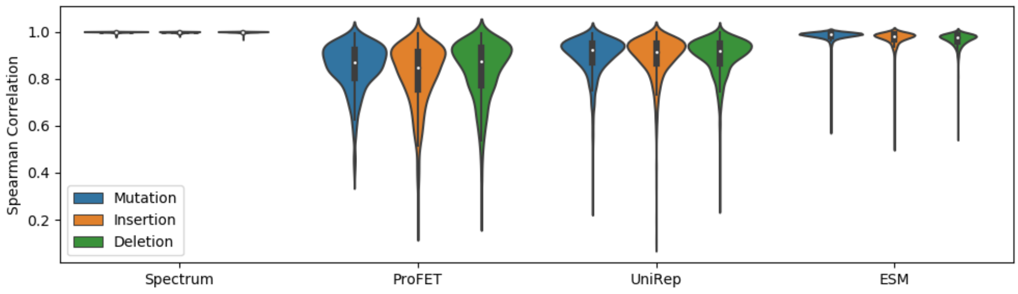


Figure S12: Distribution of Spearman correlation between string edit distance and euclidean embedding distance over 1000 sequences with different perturbations. This analysis could only be performed on the sequence-based perturbations.

### 12 P-values corresponding to MMD tests

Table S10: Empirical p-values (Monte-Carlo simulation, n=1000) to the corresponding MMD tests for three embeddings and all models of the main results (mean and standard deviation over five data splits). Recall that in the case of a good generative model we would like to confirm the null hypothesis that the two samples originate from the same distribution, and thus expect a high p-value.

| Model | Spectrum | ProFET | UniRep |
|---|---|---|---|
| Positive Control | 0.435 ± 0.360 | 0.523 ± 0.419 | 0.675 ± 0.255 |
| Negative Control | 0.000 ± 0.000 | 0.000 ± 0.000 | 0.000 ± 0.000 |
| ProteoGAN | 0.000 ± 0.000 | 0.000 ± 0.000 | 0.000 ± 0.000 |
| Predictorguided | 0.000 ± 0.000 | 0.000 ± 0.000 | 0.000 ± 0.000 |
| Non-Hierarchical | 0.000 ± 0.000 | 0.000 ± 0.000 | 0.000 ± 0.000 |
| CVAE | 0.000 ± 0.000 | 0.000 ± 0.000 | 0.000 ± 0.000 |
| OpC-ngram | 0.000 ± 0.000 | 0.000 ± 0.000 | 0.000 ± 0.000 |
| OpC-HMM | 0.000 ± 0.000 | 0.000 ± 0.000 | 0.000 ± 0.000 |
| OpL-GAN | 0.000 | 0.000 | 0.000 |
| OpL-ngram | 0.000 ± 0.000 | 0.000 ± 0.000 | 0.000 ± 0.000 |
| OpL-HMM | 0.000 ± 0.000 | 0.000 ± 0.000 | 0.000 ± 0.000 |
| ProteoGAN (100 labels) | 0.000 | 0.000 | 0.000 |
| ProteoGAN (200 labels) | 0.000 | 0.000 | 0.000 |

### 13 Kolmogorov-Smirnov statistics

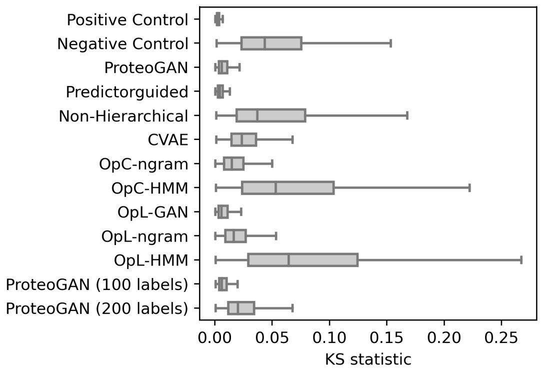


Figure S13: KS statistics over 8000 Spectrum kernel features, lower is better.

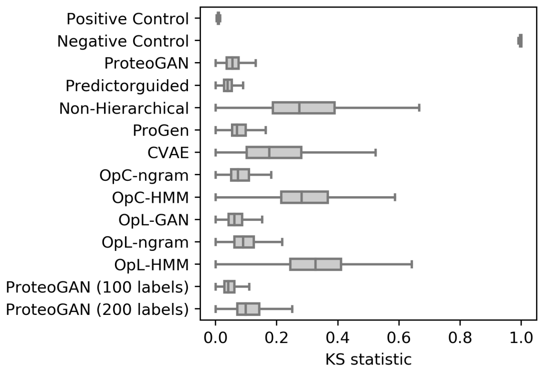


Figure S14: KS statistics over ca. 500 ProFET biological features, lower is better.

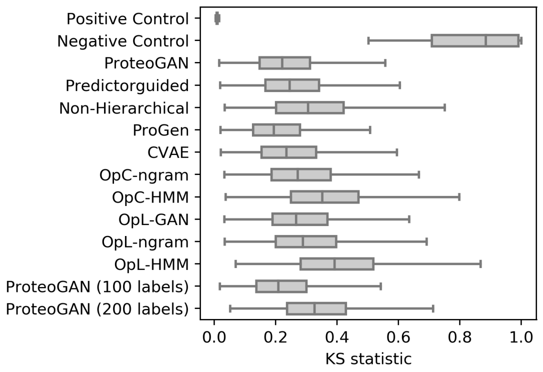


Figure S15: KS statistics over 1900 UniRep biological features, lower is better.

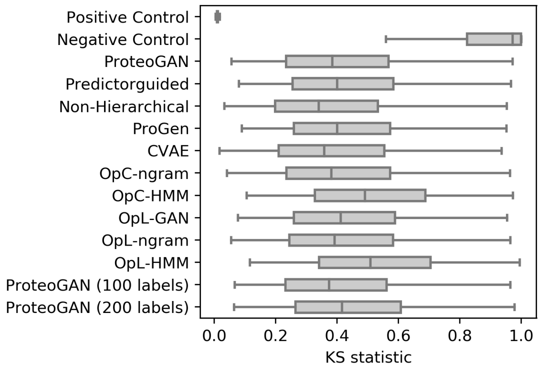


Figure S16: KS statistics over 1280 ESM biological features, lower is better.

#### 14 PCA of generated vs real sequences

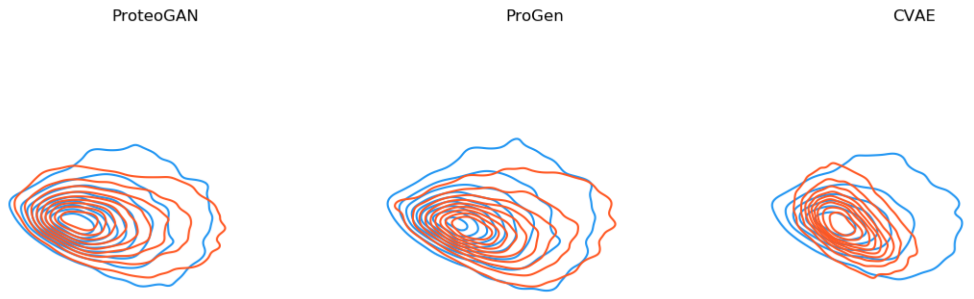


Figure S17: KDE plot on a PCA of three generative models (red) compared to the distribution of real (test set) sequences. Contours indicate density. ProteoGAN is best able to reproduce the distribution of real sequences, confirming the MMD results. Note that the two principal components only explain 1.6% of the variance, MMD statistics are a better tool to assess the two-sample problem in this case.

### 15 Model evaluation with homology-controlled test sets

Table S11: Model evaluation with homology controlled test sets and the Spectrum embedding. Sequences were allowed to have up to 90% homology with the training set. Compare main result tables.

| Model | MMD↓ | gauss. MMD↓ | MRR↑ | $MRR_B$ ↑ | ΔEntropy | ΔDistance |
|---|---|---|---|---|---|---|
| Positive Control | 0.018 ± 0.001 | 0.014 ± 0.000 | 0.769 ± 0.021 | 0.902 ± 0.023 | -0.039 ± 0.008 | 0.005 ± 0.002 |
| Negative Control | 1.014 ± 0.000 | 0.934 ± 0.000 | 0.090 ± 0.000 | 0.097 ± 0.000 | 0.688 ± 0.007 | 1.848 ± 0.002 |
| ProteoGAN | 0.045 ± 0.002 | 0.028 ± 0.001 | 0.527 ± 0.015 | 0.702 ± 0.035 | -0.052 ± 0.011 | 0.017 ± 0.004 |
| Predictorguided | 0.030 ± 0.001 | 0.021 ± 0.001 | 0.114 ± 0.004 | 0.141 ± 0.007 | -0.028 ± 0.009 | 0.007 ± 0.003 |
| Non-Hierarchical | 0.336 ± 0.118 | 0.242 ± 0.096 | 0.298 ± 0.048 | 0.421 ± 0.057 | -0.395 ± 0.179 | 0.295 ± 0.170 |
| ProGen | 0.055 | 0.034 | 0.392 | 0.518 | -0.197 | 0.042 |
| CVAE | 0.231 ± 0.077 | 0.147 ± 0.057 | 0.282 ± 0.019 | 0.398 ± 0.035 | 0.206 ± 0.026 | 0.151 ± 0.086 |
| OpC-ngram | 0.055 ± 0.002 | 0.034 ± 0.001 | 0.338 ± 0.016 | 0.408 ± 0.020 | 0.171 ± 0.006 | -0.045 ± 0.002 |
| OpC-HMM | 0.175 ± 0.003 | 0.111 ± 0.002 | 0.099 ± 0.008 | 0.145 ± 0.007 | -0.627 ± 0.016 | 0.204 ± 0.005 |
| OpL-GAN | 0.042 | 0.027 | 0.493 | 0.686 | -0.102 | 0.027 |
| OpL-ngram | 0.058 ± 0.002 | 0.036 ± 0.001 | 0.324 ± 0.011 | 0.384 ± 0.014 | 0.196 ± 0.008 | -0.047 ± 0.002 |
| OpL-HMM | 0.200 ± 0.002 | 0.129 ± 0.002 | 0.098 ± 0.009 | 0.145 ± 0.009 | -0.703 ± 0.013 | 0.249 ± 0.004 |
| ProteoGAN (100 labels) | 0.040 | 0.027 | 0.574 | 0.769 | -0.066 | 0.024 |
| ProteoGAN (200 labels) | 0.163 | 0.112 | 0.298 | 0.455 | 0.065 | 0.056 |

Table S12: Model evaluation with homology controlled test sets and the ProFET embedding. Sequences were allowed to have up to 90% homology with the training set. Compare main result tables.

| Model | MMD↓ | gauss. MMD↓ | MRR↑ | $MRR_B$ ↑ | ΔEntropy | ΔDistance |
|---|---|---|---|---|---|---|
| Positive Control | 0.066 ± 0.011 | 0.016 ± 0.001 | 0.723 ± 0.034 | 0.850 ± 0.032 | 0.002 ± 0.008 | 0.100 ± 0.096 |
| Negative Control | 11.673 ± 0.007 | 1.006 ± 0.000 | 0.090 ± 0.000 | 0.119 ± 0.000 | 7.242 ± 0.011 | 6.484 ± 0.090 |
| ProteoGAN | 0.184 ± 0.013 | 0.040 ± 0.002 | 0.509 ± 0.024 | 0.682 ± 0.029 | -0.042 ± 0.026 | -0.217 ± 0.247 |
| Predictorguided | 0.138 ± 0.006 | 0.029 ± 0.001 | 0.130 ± 0.016 | 0.166 ± 0.033 | 0.038 ± 0.024 | 0.261 ± 0.177 |
| Non-Hierarchical | 1.152 ± 0.259 | 0.142 ± 0.016 | 0.290 ± 0.044 | 0.403 ± 0.075 | -0.207 ± 0.228 | -6.896 ± 4.099 |
| ProGen | 0.322 | 0.052 | 0.372 | 0.514 | 0.227 | 0.961 |
| CVAE | 1.070 ± 0.270 | 0.099 ± 0.006 | 0.286 ± 0.014 | 0.395 ± 0.025 | -0.138 ± 0.104 | -2.730 ± 2.080 |
| OpC-ngram | 0.526 ± 0.015 | 0.049 ± 0.001 | 0.261 ± 0.009 | 0.329 ± 0.010 | 0.003 ± 0.011 | -6.693 ± 0.273 |
| OpC-HMM | 0.736 ± 0.015 | 0.204 ± 0.002 | 0.177 ± 0.011 | 0.255 ± 0.017 | 1.138 ± 0.014 | 3.048 ± 0.090 |
| OpL-GAN | 0.220 | 0.049 | 0.581 | 0.731 | 0.200 | 1.109 |
| OpL-ngram | 0.561 ± 0.017 | 0.057 ± 0.001 | 0.259 ± 0.008 | 0.319 ± 0.010 | 0.029 ± 0.010 | -6.627 ± 0.333 |
| OpL-HMM | 0.846 ± 0.010 | 0.228 ± 0.003 | 0.198 ± 0.008 | 0.261 ± 0.013 | 1.333 ± 0.006 | 3.200 ± 0.071 |
| ProteoGAN (100 labels) | 0.150 | 0.038 | 0.500 | 0.699 | -0.036 | -0.419 |
| ProteoGAN (200 labels) | 0.253 | 0.081 | 0.277 | 0.399 | 0.321 | 0.984 |

Table S13: Model evaluation with homology controlled test sets and the UniRep embedding. Sequences were allowed to have up to 90% homology with the training set. Compare main result tables.

| Model | MMD↓ | gauss. MMD↓ | MRR↑ | $MRR_B$ ↑ | ΔEntropy | ΔDistance |
|---|---|---|---|---|---|---|
| Positive Control | 0.116 ± 0.018 | 0.015 ± 0.000 | 0.766 ± 0.034 | 0.892 ± 0.020 | -0.013 ± 0.006 | 0.255 ± 0.218 |
| Negative Control | 10.627 ± 0.003 | 1.000 ± 0.000 | 0.090 ± 0.000 | 0.116 ± 0.002 | 8.134 ± 0.006 | 28.286 ± 0.206 |
| ProteoGAN | 1.509 ± 0.018 | 0.062 ± 0.003 | 0.244 ± 0.009 | 0.305 ± 0.005 | 0.852 ± 0.029 | 12.916 ± 0.758 |
| Predictorguided | 1.592 ± 0.012 | 0.069 ± 0.002 | 0.096 ± 0.006 | 0.126 ± 0.008 | 0.951 ± 0.017 | 15.309 ± 0.282 |
| Non-Hierarchical | 2.437 ± 0.471 | 0.069 ± 0.009 | 0.174 ± 0.033 | 0.229 ± 0.035 | 0.676 ± 0.163 | 5.095 ± 3.032 |
| ProGen | 1.256 | 0.055 | 0.254 | 0.311 | 0.747 | 11.288 |
| CVAE | 1.804 ± 0.205 | 0.047 ± 0.010 | 0.212 ± 0.019 | 0.267 ± 0.022 | 0.619 ± 0.108 | 7.507 ± 3.770 |
| OpC-ngram | 2.060 ± 0.007 | 0.101 ± 0.001 | 0.145 ± 0.013 | 0.179 ± 0.017 | 0.931 ± 0.007 | 16.481 ± 0.194 |
| OpC-HMM | 2.121 ± 0.019 | 0.195 ± 0.002 | 0.122 ± 0.003 | 0.155 ± 0.005 | 1.515 ± 0.012 | 19.070 ± 0.211 |
| OpL-GAN | 1.729 | 0.096 | 0.236 | 0.281 | 1.092 | 17.203 |
| OpL-ngram | 2.153 ± 0.006 | 0.119 ± 0.001 | 0.160 ± 0.012 | 0.215 ± 0.017 | 1.013 ± 0.013 | 17.729 ± 0.209 |
| OpL-HMM | 2.302 ± 0.021 | 0.244 ± 0.006 | 0.128 ± 0.007 | 0.160 ± 0.010 | 1.735 ± 0.029 | 20.209 ± 0.452 |
| ProteoGAN (100 labels) | 1.443 | 0.056 | 0.245 | 0.302 | 0.805 | 11.057 |
| ProteoGAN (200 labels) | 2.053 | 0.150 | 0.130 | 0.162 | 1.377 | 21.351 |

Table S14: Model evaluation with homology controlled test sets and the ESM embedding. Sequences were allowed to have up to 90% homology with the training set. Compare main result tables.

| Model | MMD↓ | gauss. MMD↓ | MRR↑ | $MRR_B$ ↑ | ΔEntropy | ΔDistance |
|---|---|---|---|---|---|---|
| Positive Control | 0.483 ± 0.005 | 0.014 ± 0.000 | 0.543 ± 0.030 | 0.695 ± 0.026 | -0.053 ± 0.006 | -1.143 ± 0.263 |
| Negative Control | 10.278 ± 0.001 | 1.000 ± 0.000 | 0.090 ± 0.000 | 0.113 ± 0.002 | 8.810 ± 0.005 | 25.961 ± 0.139 |
| ProteoGAN | 5.461 ± 0.027 | 0.063 ± 0.004 | 0.096 ± 0.005 | 0.112 ± 0.004 | 0.675 ± 0.035 | 16.624 ± 0.529 |
| Predictorguided | 5.395 ± 0.015 | 0.083 ± 0.003 | 0.090 ± 0.000 | 0.106 ± 0.001 | 0.795 ± 0.015 | 18.046 ± 0.223 |
| Non-Hierarchical | 5.331 ± 0.197 | 0.058 ± 0.013 | 0.093 ± 0.001 | 0.109 ± 0.002 | 0.313 ± 0.145 | 7.424 ± 5.493 |
| ProGen | 5.179 | 0.098 | 0.103 | 0.118 | 0.873 | 18.415 |
| CVAE | 5.450 ± 0.111 | 0.026 ± 0.005 | 0.094 ± 0.003 | 0.109 ± 0.003 | 0.173 ± 0.136 | 5.343 ± 4.232 |
| OpC-ngram | 5.521 ± 0.013 | 0.071 ± 0.001 | 0.093 ± 0.004 | 0.109 ± 0.004 | 0.654 ± 0.003 | 15.686 ± 0.151 |
| OpC-HMM | 6.041 ± 0.015 | 0.170 ± 0.003 | 0.091 ± 0.001 | 0.106 ± 0.002 | 1.134 ± 0.009 | 20.872 ± 0.174 |
| OpL-GAN | 5.407 | 0.097 | 0.103 | 0.118 | 0.828 | 17.835 |
| OpL-ngram | 5.521 ± 0.015 | 0.082 ± 0.001 | 0.096 ± 0.005 | 0.111 ± 0.005 | 0.699 ± 0.003 | 16.286 ± 0.232 |
| OpL-HMM | 6.221 ± 0.013 | 0.202 ± 0.002 | 0.095 ± 0.005 | 0.110 ± 0.006 | 1.228 ± 0.006 | 21.746 ± 0.142 |
| ProteoGAN (100 labels) | 5.374 | 0.061 | 0.093 | 0.108 | 0.685 | 16.305 |
| ProteoGAN (200 labels) | 5.576 | 0.104 | 0.091 | 0.106 | 0.869 | 18.179 |

Table S15: Model evaluation with homology controlled test sets and the Spectrum embedding. Sequences were allowed to have up to 70% homology with the training set. Compare main result tables.

| Model | MMD↓ | gauss. MMD↓ | MRR↑ | $MRR_B$ ↑ | ΔEntropy | ΔDistance |
|---|---|---|---|---|---|---|
| Positive Control | 0.026 ± 0.001 | 0.020 ± 0.000 | 0.545 ± 0.033 | 0.738 ± 0.018 | -0.058 ± 0.009 | 0.006 ± 0.002 |
| Negative Control | 1.016 ± 0.001 | 0.935 ± 0.000 | 0.090 ± 0.000 | 0.098 ± 0.000 | 0.671 ± 0.007 | 1.849 ± 0.002 |
| ProteoGAN | 0.049 ± 0.002 | 0.031 ± 0.001 | 0.416 ± 0.032 | 0.580 ± 0.034 | -0.071 ± 0.010 | 0.018 ± 0.004 |
| Predictorguided | 0.035 ± 0.001 | 0.024 ± 0.001 | 0.133 ± 0.006 | 0.169 ± 0.010 | -0.047 ± 0.006 | 0.008 ± 0.003 |
| Non-Hierarchical | 0.337 ± 0.117 | 0.243 ± 0.096 | 0.293 ± 0.041 | 0.407 ± 0.054 | -0.417 ± 0.181 | 0.296 ± 0.171 |
| ProGen | 0.058 | 0.036 | 0.329 | 0.480 | -0.209 | 0.042 |
| CVAE | 0.235 ± 0.077 | 0.149 ± 0.057 | 0.256 ± 0.017 | 0.368 ± 0.013 | 0.187 ± 0.031 | 0.152 ± 0.086 |
| OpC-ngram | 0.056 ± 0.002 | 0.035 ± 0.001 | 0.298 ± 0.006 | 0.378 ± 0.009 | 0.156 ± 0.008 | -0.044 ± 0.002 |
| OpC-HMM | 0.175 ± 0.003 | 0.111 ± 0.002 | 0.110 ± 0.008 | 0.158 ± 0.010 | -0.655 ± 0.015 | 0.206 ± 0.005 |
| OpL-GAN | 0.044 | 0.029 | 0.368 | 0.533 | -0.114 | 0.027 |
| OpL-ngram | 0.059 ± 0.002 | 0.036 ± 0.001 | 0.296 ± 0.015 | 0.378 ± 0.025 | 0.181 ± 0.008 | -0.046 ± 0.002 |
| OpL-HMM | 0.201 ± 0.002 | 0.129 ± 0.001 | 0.102 ± 0.010 | 0.150 ± 0.011 | -0.732 ± 0.014 | 0.251 ± 0.003 |
| ProteoGAN (100 labels) | 0.046 | 0.031 | 0.511 | 0.681 | -0.078 | 0.024 |
| ProteoGAN (200 labels) | 0.163 | 0.113 | 0.280 | 0.417 | 0.053 | 0.056 |

Table S16: Model evaluation with homology controlled test sets and the ProFET embedding. Sequences were allowed to have up to 70% homology with the training set. Compare main result tables.

| Model | MMD↓ | gauss. MMD↓ | MRR↑ | $MRR_B$ ↑ | ΔEntropy | ΔDistance |
|---|---|---|---|---|---|---|
| Positive Control | 0.092 ± 0.011 | 0.021 ± 0.001 | 0.668 ± 0.051 | 0.802 ± 0.046 | -0.012 ± 0.009 | 0.292 ± 0.140 |
| Negative Control | 11.675 ± 0.011 | 1.006 ± 0.000 | 0.090 ± 0.000 | 0.119 ± 0.000 | 7.267 ± 0.027 | 6.676 ± 0.136 |
| ProteoGAN | 0.210 ± 0.013 | 0.043 ± 0.002 | 0.475 ± 0.020 | 0.627 ± 0.030 | -0.058 ± 0.028 | -0.025 ± 0.298 |
| Predictorguided | 0.140 ± 0.005 | 0.032 ± 0.001 | 0.127 ± 0.027 | 0.165 ± 0.025 | 0.023 ± 0.025 | 0.453 ± 0.206 |
| Non-Hierarchical | 1.159 ± 0.254 | 0.141 ± 0.017 | 0.290 ± 0.044 | 0.392 ± 0.066 | -0.219 ± 0.224 | -6.704 ± 4.068 |
| ProGen | 0.323 | 0.054 | 0.300 | 0.428 | 0.212 | 1.131 |
| CVAE | 1.075 ± 0.272 | 0.098 ± 0.006 | 0.263 ± 0.029 | 0.348 ± 0.052 | -0.152 ± 0.102 | -2.538 ± 2.059 |
| OpC-ngram | 0.510 ± 0.019 | 0.051 ± 0.001 | 0.239 ± 0.009 | 0.294 ± 0.011 | -0.010 ± 0.011 | -6.501 ± 0.294 |
| OpC-HMM | 0.732 ± 0.016 | 0.204 ± 0.002 | 0.164 ± 0.011 | 0.240 ± 0.013 | 1.120 ± 0.017 | 3.239 ± 0.136 |
| OpL-GAN | 0.216 | 0.051 | 0.574 | 0.720 | 0.184 | 1.280 |
| OpL-ngram | 0.541 ± 0.021 | 0.057 ± 0.001 | 0.251 ± 0.008 | 0.311 ± 0.012 | 0.017 ± 0.011 | -6.436 ± 0.370 |
| OpL-HMM | 0.844 ± 0.011 | 0.228 ± 0.002 | 0.183 ± 0.010 | 0.255 ± 0.030 | 1.315 ± 0.008 | 3.392 ± 0.119 |
| ProteoGAN (100 labels) | 0.178 | 0.040 | 0.476 | 0.641 | -0.051 | -0.248 |
| ProteoGAN (200 labels) | 0.239 | 0.081 | 0.276 | 0.386 | 0.304 | 1.154 |

Table S17: Model evaluation with homology controlled test sets and the UniRep embedding. Sequences were allowed to have up to 70% homology with the training set. Compare main result tables.

| Model | MMD↓ | gauss. MMD↓ | MRR↑ | $MRR_B$ ↑ | ΔEntropy | ΔDistance |
|---|---|---|---|---|---|---|
| Positive Control | 0.163 ± 0.008 | 0.019 ± 0.000 | 0.759 ± 0.019 | 0.884 ± 0.024 | -0.059 ± 0.008 | 0.383 ± 0.344 |
| Negative Control | 10.628 ± 0.004 | 1.000 ± 0.000 | 0.090 ± 0.000 | 0.112 ± 0.003 | 8.127 ± 0.004 | 28.414 ± 0.355 |
| ProteoGAN | 1.497 ± 0.018 | 0.062 ± 0.003 | 0.228 ± 0.012 | 0.288 ± 0.016 | 0.803 ± 0.030 | 13.044 ± 0.874 |
| Predictorguided | 1.574 ± 0.014 | 0.069 ± 0.002 | 0.099 ± 0.006 | 0.125 ± 0.007 | 0.903 ± 0.019 | 15.437 ± 0.413 |
| Non-Hierarchical | 2.439 ± 0.468 | 0.069 ± 0.010 | 0.171 ± 0.029 | 0.228 ± 0.033 | 0.629 ± 0.159 | 5.223 ± 3.010 |
| ProGen | 1.261 | 0.055 | 0.239 | 0.312 | 0.706 | 11.634 |
| CVAE | 1.812 ± 0.201 | 0.046 ± 0.010 | 0.209 ± 0.026 | 0.259 ± 0.024 | 0.570 ± 0.111 | 7.634 ± 3.876 |
| OpC-ngram | 2.047 ± 0.016 | 0.101 ± 0.001 | 0.137 ± 0.010 | 0.164 ± 0.013 | 0.886 ± 0.007 | 16.608 ± 0.325 |
| OpC-HMM | 2.094 ± 0.023 | 0.195 ± 0.002 | 0.122 ± 0.009 | 0.148 ± 0.011 | 1.465 ± 0.013 | 19.197 ± 0.361 |
| OpL-GAN | 1.725 | 0.096 | 0.212 | 0.274 | 1.049 | 17.548 |
| OpL-ngram | 2.137 ± 0.017 | 0.119 ± 0.001 | 0.159 ± 0.012 | 0.203 ± 0.009 | 0.967 ± 0.013 | 17.857 ± 0.338 |
| OpL-HMM | 2.275 ± 0.026 | 0.244 ± 0.006 | 0.123 ± 0.007 | 0.148 ± 0.010 | 1.684 ± 0.030 | 20.337 ± 0.532 |
| ProteoGAN (100 labels) | 1.452 | 0.055 | 0.235 | 0.292 | 0.762 | 11.403 |
| ProteoGAN (200 labels) | 2.052 | 0.150 | 0.127 | 0.161 | 1.334 | 21.697 |

Table S18: Model evaluation with homology controlled test sets and the ESM embedding. Sequences were allowed to have up to 70% homology with the training set. Compare main result tables.

| Model | MMD↓ | gauss. MMD↓ | MRR↑ | $MRR_B$ ↑ | ΔEntropy | ΔDistance |
|---|---|---|---|---|---|---|
| Positive Control | 0.911 ± 0.009 | 0.018 ± 0.000 | 0.415 ± 0.010 | 0.585 ± 0.014 | -0.138 ± 0.006 | -2.271 ± 0.278 |
| Negative Control | 10.261 ± 0.003 | 1.000 ± 0.000 | 0.090 ± 0.000 | 0.113 ± 0.005 | 8.787 ± 0.003 | 24.833 ± 0.165 |
| ProteoGAN | 5.207 ± 0.025 | 0.064 ± 0.004 | 0.094 ± 0.005 | 0.110 ± 0.005 | 0.585 ± 0.035 | 15.496 ± 0.558 |
| Predictorguided | 5.140 ± 0.023 | 0.083 ± 0.003 | 0.090 ± 0.001 | 0.107 ± 0.001 | 0.705 ± 0.014 | 16.918 ± 0.177 |
| Non-Hierarchical | 5.088 ± 0.201 | 0.059 ± 0.012 | 0.093 ± 0.001 | 0.109 ± 0.001 | 0.225 ± 0.141 | 6.296 ± 5.492 |
| ProGen | 4.919 | 0.098 | 0.091 | 0.107 | 0.783 | 17.174 |
| CVAE | 5.217 ± 0.120 | 0.027 ± 0.004 | 0.095 ± 0.003 | 0.111 ± 0.003 | 0.086 ± 0.135 | 4.215 ± 4.228 |
| OpC-ngram | 5.279 ± 0.017 | 0.072 ± 0.001 | 0.091 ± 0.001 | 0.107 ± 0.001 | 0.571 ± 0.005 | 14.558 ± 0.167 |
| OpC-HMM | 5.793 ± 0.016 | 0.170 ± 0.003 | 0.091 ± 0.001 | 0.107 ± 0.001 | 1.042 ± 0.009 | 19.744 ± 0.197 |
| OpL-GAN | 5.153 | 0.098 | 0.091 | 0.107 | 0.736 | 16.594 |
| OpL-ngram | 5.277 ± 0.019 | 0.082 ± 0.001 | 0.091 ± 0.001 | 0.108 ± 0.001 | 0.613 ± 0.004 | 15.159 ± 0.289 |
| OpL-HMM | 5.975 ± 0.014 | 0.202 ± 0.002 | 0.090 ± 0.001 | 0.106 ± 0.001 | 1.136 ± 0.005 | 20.618 ± 0.159 |
| ProteoGAN (100 labels) | 5.118 | 0.061 | 0.091 | 0.107 | 0.594 | 15.064 |
| ProteoGAN (200 labels) | 5.324 | 0.104 | 0.090 | 0.106 | 0.778 | 16.938 |

Table S19: Model evaluation with homology controlled test sets and the Spectrum embedding. Sequences were allowed to have up to 50% homology with the training set. Compare main result tables.

| Model | MMD↓ | gauss. MMD↓ | MRR↑ | $MRR_B$ ↑ | ΔEntropy | ΔDistance |
|---|---|---|---|---|---|---|
| Positive Control | 0.038 ± 0.000 | 0.028 ± 0.000 | 0.299 ± 0.013 | 0.447 ± 0.016 | -0.084 ± 0.008 | 0.006 ± 0.003 |
| Negative Control | 1.018 ± 0.001 | 0.936 ± 0.000 | 0.090 ± 0.000 | 0.099 ± 0.001 | 0.645 ± 0.007 | 1.849 ± 0.002 |
| ProteoGAN | 0.056 ± 0.003 | 0.037 ± 0.001 | 0.310 ± 0.032 | 0.426 ± 0.041 | -0.099 ± 0.011 | 0.019 ± 0.004 |
| Predictorguided | 0.042 ± 0.002 | 0.030 ± 0.001 | 0.114 ± 0.007 | 0.150 ± 0.014 | -0.074 ± 0.009 | 0.008 ± 0.004 |
| Non-Hierarchical | 0.338 ± 0.117 | 0.243 ± 0.095 | 0.276 ± 0.045 | 0.405 ± 0.057 | -0.449 ± 0.188 | 0.296 ± 0.169 |
| ProGen | 0.063 | 0.041 | 0.223 | 0.352 | -0.243 | 0.043 |
| CVAE | 0.239 ± 0.076 | 0.153 ± 0.056 | 0.192 ± 0.018 | 0.299 ± 0.022 | 0.160 ± 0.030 | 0.152 ± 0.085 |
| OpC-ngram | 0.060 ± 0.002 | 0.040 ± 0.001 | 0.236 ± 0.008 | 0.321 ± 0.015 | 0.136 ± 0.007 | -0.044 ± 0.003 |
| OpC-HMM | 0.176 ± 0.003 | 0.112 ± 0.002 | 0.103 ± 0.013 | 0.157 ± 0.025 | -0.695 ± 0.019 | 0.206 ± 0.006 |
| OpL-GAN | 0.051 | 0.035 | 0.235 | 0.359 | -0.147 | 0.028 |
| OpL-ngram | 0.061 ± 0.002 | 0.040 ± 0.001 | 0.272 ± 0.017 | 0.371 ± 0.020 | 0.161 ± 0.009 | -0.046 ± 0.003 |
| OpL-HMM | 0.201 ± 0.002 | 0.130 ± 0.002 | 0.099 ± 0.011 | 0.153 ± 0.025 | -0.776 ± 0.016 | 0.251 ± 0.004 |
| ProteoGAN (100 labels) | 0.056 | 0.038 | 0.377 | 0.486 | -0.111 | 0.025 |
| ProteoGAN (200 labels) | 0.165 | 0.115 | 0.197 | 0.313 | 0.020 | 0.057 |

Table S20: Model evaluation with homology controlled test sets and the ProFET embedding. Sequences were allowed to have up to 50% homology with the training set. Compare main result tables.

| Model | MMD↓ | gauss. MMD↓ | MRR↑ | $MRR_B$ ↑ | ΔEntropy | ΔDistance |
|---|---|---|---|---|---|---|
| Positive Control | 0.136 ± 0.010 | 0.032 ± 0.001 | 0.547 ± 0.024 | 0.674 ± 0.012 | -0.054 ± 0.016 | 0.752 ± 0.263 |
| Negative Control | 11.668 ± 0.014 | 1.005 ± 0.000 | 0.090 ± 0.000 | 0.119 ± 0.000 | 7.284 ± 0.030 | 7.136 ± 0.259 |
| ProteoGAN | 0.244 ± 0.021 | 0.047 ± 0.003 | 0.445 ± 0.053 | 0.602 ± 0.062 | -0.107 ± 0.035 | 0.434 ± 0.397 |
| Predictorguided | 0.158 ± 0.013 | 0.038 ± 0.002 | 0.130 ± 0.021 | 0.171 ± 0.035 | -0.022 ± 0.030 | 0.913 ± 0.338 |
| Non-Hierarchical | 1.163 ± 0.250 | 0.139 ± 0.017 | 0.265 ± 0.044 | 0.375 ± 0.077 | -0.260 ± 0.213 | -6.244 ± 4.028 |
| ProGen | 0.339 | 0.060 | 0.292 | 0.434 | 0.171 | 1.621 |
| CVAE | 1.083 ± 0.277 | 0.096 ± 0.005 | 0.246 ± 0.035 | 0.343 ± 0.051 | -0.196 ± 0.100 | -2.078 ± 1.973 |
| OpC-ngram | 0.493 ± 0.020 | 0.055 ± 0.001 | 0.229 ± 0.013 | 0.281 ± 0.017 | -0.051 ± 0.015 | -6.041 ± 0.413 |
| OpC-HMM | 0.730 ± 0.021 | 0.206 ± 0.002 | 0.170 ± 0.017 | 0.246 ± 0.025 | 1.062 ± 0.025 | 3.699 ± 0.259 |
| OpL-GAN | 0.231 | 0.057 | 0.438 | 0.573 | 0.141 | 1.769 |
| OpL-ngram | 0.519 ± 0.025 | 0.061 ± 0.001 | 0.236 ± 0.032 | 0.287 ± 0.036 | -0.024 ± 0.017 | -5.976 ± 0.392 |
| OpL-HMM | 0.841 ± 0.016 | 0.229 ± 0.003 | 0.168 ± 0.027 | 0.227 ± 0.043 | 1.258 ± 0.016 | 3.851 ± 0.238 |
| ProteoGAN (100 labels) | 0.230 | 0.045 | 0.423 | 0.576 | -0.095 | 0.241 |
| ProteoGAN (200 labels) | 0.243 | 0.084 | 0.252 | 0.339 | 0.257 | 1.644 |

Table S21: Model evaluation with homology controlled test sets and the UniRep embedding. Sequences were allowed to have up to 50% homology with the training set. Compare main result tables.

| Model | MMD↓ | gauss. MMD↓ | MRR↑ | $MRR_B$ ↑ | ΔEntropy | ΔDistance |
|---|---|---|---|---|---|---|
| Positive Control | 0.292 ± 0.017 | 0.029 ± 0.000 | 0.725 ± 0.032 | 0.870 ± 0.022 | -0.202 ± 0.016 | 0.577 ± 0.387 |
| Negative Control | 10.604 ± 0.004 | 1.000 ± 0.000 | 0.090 ± 0.000 | 0.106 ± 0.003 | 8.063 ± 0.012 | 28.608 ± 0.395 |
| ProteoGAN | 1.402 ± 0.020 | 0.062 ± 0.003 | 0.240 ± 0.018 | 0.289 ± 0.018 | 0.645 ± 0.038 | 13.238 ± 0.968 |
| Predictorguided | 1.474 ± 0.012 | 0.069 ± 0.003 | 0.095 ± 0.006 | 0.117 ± 0.006 | 0.749 ± 0.024 | 15.631 ± 0.314 |
| Non-Hierarchical | 2.369 ± 0.464 | 0.069 ± 0.009 | 0.176 ± 0.028 | 0.229 ± 0.028 | 0.479 ± 0.144 | 5.417 ± 2.844 |
| ProGen | 1.150 | 0.055 | 0.257 | 0.328 | 0.538 | 11.308 |
| CVAE | 1.746 ± 0.214 | 0.047 ± 0.010 | 0.197 ± 0.013 | 0.246 ± 0.015 | 0.416 ± 0.098 | 7.829 ± 3.602 |
| OpC-ngram | 1.976 ± 0.008 | 0.101 ± 0.001 | 0.134 ± 0.009 | 0.162 ± 0.011 | 0.740 ± 0.019 | 16.803 ± 0.342 |
| OpC-HMM | 1.981 ± 0.025 | 0.195 ± 0.002 | 0.118 ± 0.010 | 0.142 ± 0.009 | 1.305 ± 0.020 | 19.392 ± 0.433 |
| OpL-GAN | 1.589 | 0.095 | 0.203 | 0.262 | 0.874 | 17.222 |
| OpL-ngram | 2.065 ± 0.008 | 0.119 ± 0.002 | 0.149 ± 0.010 | 0.196 ± 0.011 | 0.821 ± 0.021 | 18.052 ± 0.444 |
| OpL-HMM | 2.162 ± 0.016 | 0.244 ± 0.006 | 0.118 ± 0.013 | 0.142 ± 0.013 | 1.524 ± 0.028 | 20.531 ± 0.630 |
| ProteoGAN (100 labels) | 1.334 | 0.055 | 0.250 | 0.308 | 0.588 | 11.076 |
| ProteoGAN (200 labels) | 1.930 | 0.149 | 0.105 | 0.125 | 1.157 | 21.371 |

Table S22: Model evaluation with homology controlled test sets and the ESM embedding. Sequences were allowed to have up to 50% homology with the training set. Compare main result tables.

| Model | MMD↓ | gauss. MMD↓ | MRR↑ | $MRR_B$ ↑ | ΔEntropy | ΔDistance |
|---|---|---|---|---|---|---|
| Positive Control | 1.386 ± 0.029 | 0.024 ± 0.000 | 0.287 ± 0.027 | 0.420 ± 0.029 | -0.304 ± 0.011 | -3.177 ± 0.385 |
| Negative Control | 10.214 ± 0.007 | 1.000 ± 0.000 | 0.090 ± 0.000 | 0.114 ± 0.001 | 8.708 ± 0.009 | 23.928 ± 0.312 |
| ProteoGAN | 4.797 ± 0.040 | 0.066 ± 0.004 | 0.092 ± 0.001 | 0.107 ± 0.001 | 0.398 ± 0.040 | 14.590 ± 0.743 |
| Predictorguided | 4.727 ± 0.032 | 0.085 ± 0.003 | 0.090 ± 0.000 | 0.106 ± 0.001 | 0.517 ± 0.022 | 16.012 ± 0.364 |
| Non-Hierarchical | 4.722 ± 0.208 | 0.061 ± 0.012 | 0.092 ± 0.001 | 0.108 ± 0.001 | 0.045 ± 0.125 | 5.390 ± 5.430 |
| ProGen | 4.484 | 0.100 | 0.092 | 0.107 | 0.602 | 16.091 |
| CVAE | 4.861 ± 0.160 | 0.032 ± 0.004 | 0.092 ± 0.001 | 0.109 ± 0.002 | -0.092 ± 0.126 | 3.309 ± 4.098 |
| OpC-ngram | 4.876 ± 0.032 | 0.074 ± 0.001 | 0.091 ± 0.001 | 0.107 ± 0.001 | 0.407 ± 0.010 | 13.652 ± 0.371 |
| OpC-HMM | 5.389 ± 0.038 | 0.171 ± 0.003 | 0.091 ± 0.001 | 0.106 ± 0.001 | 0.848 ± 0.017 | 18.838 ± 0.350 |
| OpL-GAN | 4.726 | 0.099 | 0.092 | 0.107 | 0.549 | 15.510 |
| OpL-ngram | 4.873 ± 0.031 | 0.084 ± 0.001 | 0.091 ± 0.001 | 0.107 ± 0.001 | 0.438 ± 0.009 | 14.253 ± 0.436 |
| OpL-HMM | 5.573 ± 0.033 | 0.203 ± 0.002 | 0.091 ± 0.001 | 0.106 ± 0.000 | 0.941 ± 0.014 | 19.712 ± 0.315 |
| ProteoGAN (100 labels) | 4.689 | 0.063 | 0.092 | 0.107 | 0.404 | 13.981 |
| ProteoGAN (200 labels) | 4.899 | 0.106 | 0.090 | 0.105 | 0.586 | 15.854 |

### 16 Comparison with random mutagenesis

Table S23: Comparison of MMD achieved by ProteoGAN and MMD of a natural set of proteins with increasing amounts of random mutation, simulating experimental random mutagenesis with different mutation rates. MMD values are computed with the Spectrum embedding. Indicated are the sequence identities of the mutated sequences with their original. Random mutagenesis with 90% sequence identity achieved similar MMD values to ProteoGAN, although ProteoGAN generates sequences with only 56% sequence identity. ProteoGAN can thus generate sequences that are four to five times more novel (contain more mutations) than random mutagenesis, at the same deviation from the protein distribution.

| Sequence Identity | MMD |
|---|---|
| 100% | 0.011 |
| 99% | 0.011 |
| 98% | 0.014 |
| 97% | 0.017 |
| 96% | 0.021 |
| 95% | 0.024 |
| 94% | 0.029 |
| 93% | 0.033 |
| 92% | 0.037 |
| 91% | 0.041 |
| 90% | 0.045 |
| ProteoGAN (56%) | 0.044 |

### 17 OOD evaluation

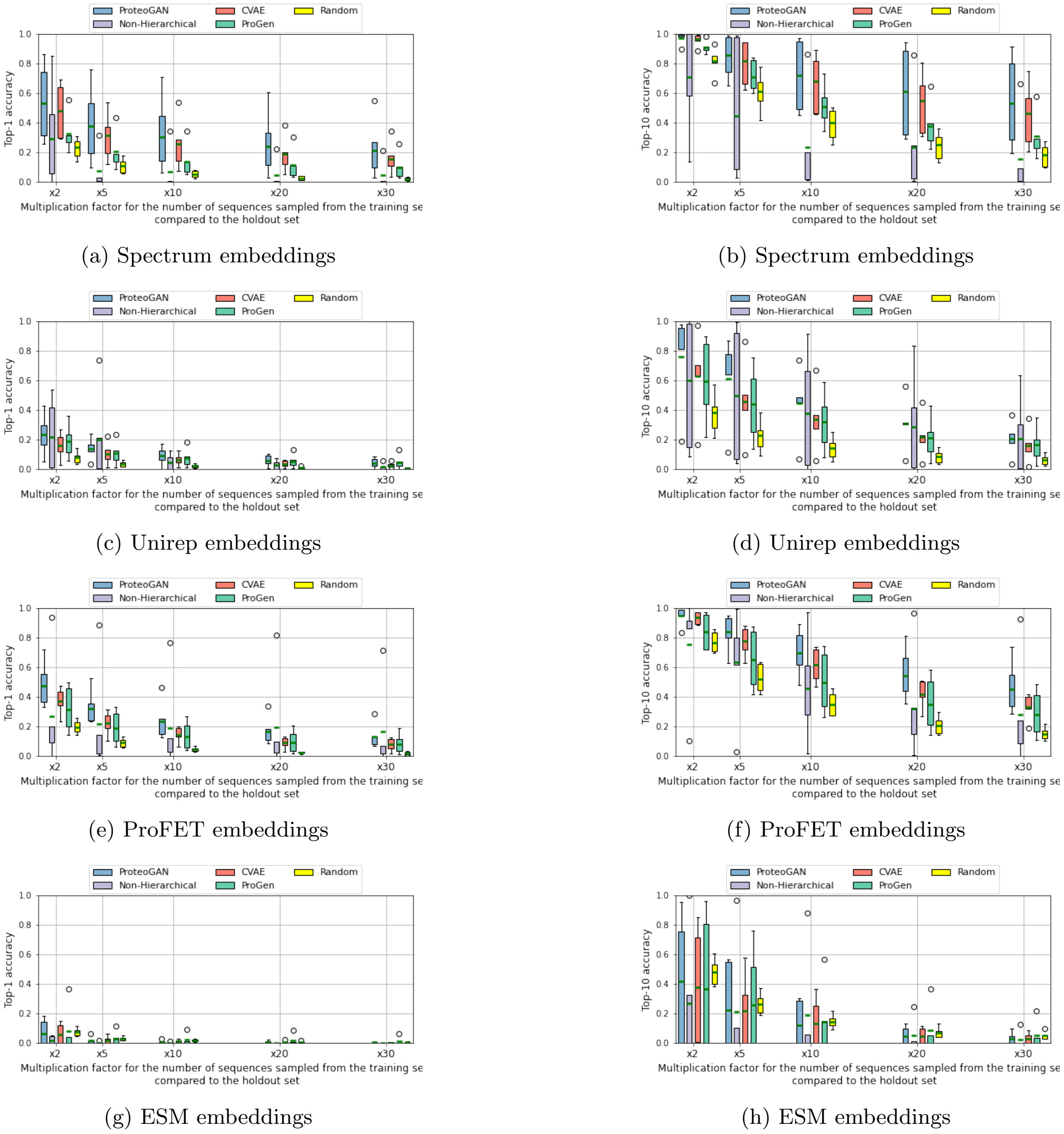


(a) Spectrum embeddings

(b) Spectrum embeddings

(c) Unirep embeddings

(d) Unirep embeddings

(e) ProFET embeddings

(f) ProFET embeddings

(g) ESM embeddings

(h) ESM embeddings

Figure S18: Top-1 and Top-10 accuracies for ProteoGAN, *Non-Hierachical*, CVAE, ProGen and a Random baseline, which consists in sampling real sequences from the training set. Distances between generated sequences and real sequences are calculated in the embedding spaces, either Spectrum, Unirep, ProFET or ESM.

### 18 Multiple Sequence Alignments of some generated sequences

We computed Multiple Sequence Alignments (MSAs) for three models (ProteoGAN, CVAE, ProGen) on a few generated sequences conditioned on the same label set (labels were taken from the holdout sets, compare Table S2). Some excerpts are replicated below, full data can be found at `https://github.com/timkucera/proteogan/tree/master/paper/supplemental_data`.

Generally, ProteoGAN and ProGen showed few conserved residues in their generated sequences. CVAE showed a block-aligned structure with more conservation. Sequences between models are very dissimilar with few aligned positions, however some short blocks of aligned residues exist between models.

Representative part of MSAs for each model independently:

```
ProteoGAN:B --------------APLAYPRPGPLIAFLAPADSFQQSQPSIDLELVTHSGL--------
ProteoGAN:B TCRDSMEKDESREYGPSLYSDFGVINKFSQNYRKSTRPV--------IKSSSKKVKLGVS
ProteoGAN:B ----------------YYCPQKGPVDKFVKEVVKRYTPAQSRRTEANDT-----------
ProteoGAN:B --------------------GGKPLDAFASRNVDD--------IEAKT------------
ProteoGAN:B --------------NMATYMANENIPAFLMRSVSFQEGMSSLEFGAGVSELTKRIRLRKT

ProGen:B    ------------------------------------------------------------
ProGen:B    RKIEANLKRPKEAVESFFNPSKGIFMYGIFEPMNQQWINGFLSEQTCYSQAAFKDKKTLS
ProGen:B    D--GCTYKRTNFVNKSY------------IYTAQGR--------------------RKRS
ProGen:B    K--GCPYESFYSK--------------------CSA--------------------LKQN
ProGen:B    VGALCNLMRQHIGSPNR------------LYLTEAKWCPGVMSVITCFLDVNCIHCKLQW

CVAE:B      MAKDKAEKELTKPKRKSKRRLGEREKPRAVDLIRLRHKDRQLSKVIQSRKDKRKVKAIRG
CVAE:B      MASEGAQTDSAKKAPESKLVLGARHVEKGLTLSVKVSLLERLSGGKVAATDKKYTKV---
CVAE:B      MQKDHNQKPGDKRSPVAESVLPNREHTEKPPPGVGVNRDQRRSKRSEAVRADRRREKSSV
CVAE:B      MAKGEAEKPETKDKGESGGVLGDAEKTRDGDLIRKSHLDRKLEKVIQSSNAKHKRKAKDE
CVAE:B      MAKERVQTTLAVLGSEAAKVLAAAGAEKEKSGSLDVVLDNRLSKVKVSMRKLVKVLVSDV
```

Representative part of MSAs between all three models:

```
ProteoGAN:B -------QS-QPSIDLELVTHSGLN--RARSLRMGMRR---SGGPGQCSLD---------
ProteoGAN:B PDGLNLPDN-VNILSASLTFHCID----LCEEVLGEAK---KQGKEN-------------
ProteoGAN:B ------PQV-VQAIGRQQLEDLNPV--QKRNKIVSQSG---KP-----------------
ProteoGAN:B ----------------------------IGEPVLEEEK---DGEKSTELRK------RSQ
ProteoGAN:B ------------RLA-------------AKRRIMVKRS---KPRIPPERAE------RFD
CVAE:B      ----------------------------KSKRRLGERE---KPRAVDLIRL---------
CVAE:B      ----------------------------ESKLVLGARH---VEKGLTLSVK---------
CVAE:B      ----------------------------VAESVLPNRE---HTEKPPPGVG---------
CVAE:B      ----------------------------ESGGVLGDAE---KTRDGDLIRK---------
CVAE:B      ----------------------------EAAKVLAAAG---AEKEKSGSLD---------
ProGen:B    ------------------------------------------DCGAEIA-----------
ProGen:B    ----------MDNLNSLILFKLKNK--QQRNELYGEFM----------------------
ProGen:B    QDGLRYHQEAVSANGDIILWNANKAAAELRQEVLDKVTAYKVGSGITVVVYAATHDDAVR
ProGen:B    ---MSQNQY-HRTFQALQLQDMER----VQEELVSE----------QPRWE------TLR
ProGen:B    -------WI---FLRALVVCQVGPT--CSADSLLRDF----DGSEAHESWQ------RL-
```

### 19 Structure predictions of ProteoGAN generated proteins

We obtained structure predictions for a small sample ($n = 50$) of generated sequences with trRosetta [16] (max_iter= 200). Out of three predictions for each sequence we selected the best by choosing the one with minimal energy as predicted by the Rosetta all-atom energy function [1]. We then measure how much the designed proteins resemble natural folds by aligning [2] their structures to the Protein Data Bank (PDB). The structural similarities are reported as TM-score [17], where higher is better. Some exemplary structure alignments are shown in Figure S19, the average TM-score was $0.60 \pm 0.09$.

We note however that this prediction pipeline was not tested on generated sequences before and that it is not guaranteed to work as expected with natural proteins.

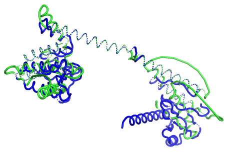

(a) TM-Score = 0.65

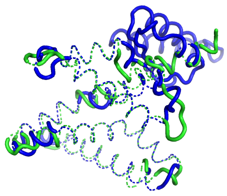

(b) TM-Score = 0.79

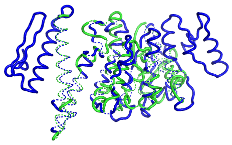

(c) TM-Score = 0.65

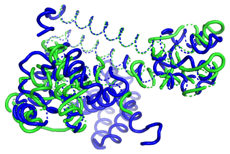

(d) TM-Score = 0.68

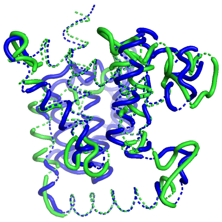

(e) TM-Score = 0.72

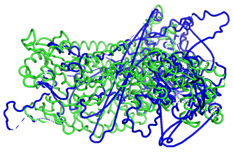

(f) TM-Score = 0.52

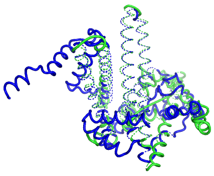

(g) TM-Score = 0.71

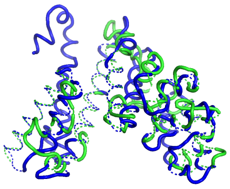

(h) TM-Score = 0.69

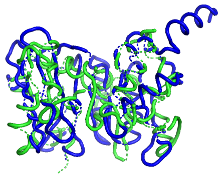

(i) TM-Score = 0.61

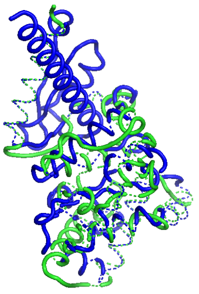

(j) TM-Score = 0.65

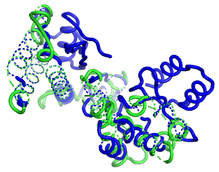

(k) TM-Score = 0.69

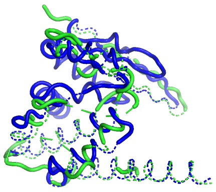

(l) TM-Score = 0.72

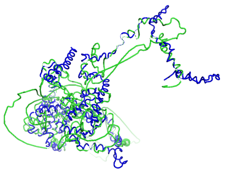

(m) TM-Score = 0.54

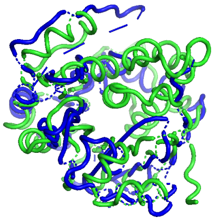

(n) TM-Score = 0.61

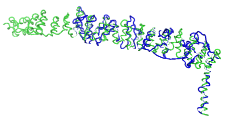

(o) TM-Score = 0.58

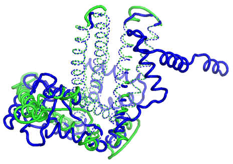

(p) TM-Score = 0.72

Figure S19: Some exemplary ProteoGAN generated structures (blue) in comparison to their structurally closest homolog in PDB (green).
